# The Protein Disulfide Isomerase P4HB/PDIA1 Modulates PrP^C^ Levels and Prion Replication

**DOI:** 10.64898/2025.12.01.691611

**Authors:** Genki Amano, Hamza Arshad, Zeel Patel, Gerold Schmitt-Ulms, Joel C. Watts

## Abstract

Prions are misfolded, self-propagating versions of the cellular prion protein (PrP^C^) that cause invariably fatal transmissible neurodegenerative diseases in humans and animals. Little is known about how prions replicate in the brain, including whether other proteins participate in prion replication *in vivo*. Several members of the protein disulfide isomerase family have been shown to reside in close spatial proximity to PrP^C^ in cells and mice, implying that they could be involved in prion biogenesis. Here, we show that stable knock-down of the protein disulfide isomerase P4HB (also called PDIA1) in prion-susceptible CAD5 cells reduces PrP^C^ levels and results in lower levels of protease-resistant PrP (PrP^res^), a marker of infectious prions, following infection with two different prion strains. Partial reduction of P4HB activity using the P4HB-selective inhibitor KSC-34 also decreases PrP^C^ levels in uninfected CAD5 cells whereas treatment of prion-infected CAD5 cells with KSC-34 results in higher levels of PrP^res^. A proportion of P4HB reaches the cell surface where PrP^C^ is located, and a secreted P4HB variant increases PrP^res^ levels in cells. Collectively, these results suggest that P4HB facilitates prion replication in cells by stabilizing PrP^C^ and potentially acting as a chaperone that directly modulates prion conversion. Thus, targeting P4HB during prion disease may have therapeutic benefit.

## Introduction

Prions are pathogenic and infectious protein aggregates that cause a variety of fatal neurodegenerative conditions in both humans and animals collectively referred to as prion diseases [1–3]. Prion diseases such as Creutzfeldt–Jakob disease (CJD) in humans are caused by the accumulation of misfolded prion protein (PrP) in cells of the central nervous system. Cellular PrP (PrP^C^) has a predominantly α-helical structure and is attached to the outer leaflet of the cell membrane of neurons and other central nervous system cells [4, 5]. PrP^C^ biosynthesis involves transit through the secretory pathway where a series of post-translational modifications occur in the endoplasmic reticulum (ER) and Golgi apparatus, including the formation of an internal disulfide bond between cysteine residues 178 and 213 in mouse PrP, the addition of up to two N-linked glycans, as well as attachment of a glycosylphosphatidylinositol (GPI) anchor at the C-terminal end of the protein. During prion disease, PrP^C^ misfolds to form β-sheet-rich aggregates termed PrP^Sc^ that are insoluble in non-ionic detergents and partially resistant to digestion with proteinase K (PK) [6–9]. Prions can exist as distinct strains, each of which is associated with a unique PrP^Sc^ structure [10, 11]. PrP^Sc^ is thought to propagate via a self-templating mechanism in which PrP^Sc^ aggregates convert PrP^C^ into additional PrP^Sc^, enabling prion replication and spreading of PrP^Sc^ within and to the central nervous system, ultimately leading to neuronal vacuolation and neurodegeneration. There are currently no therapeutics available to treat prion disease. While PrP^C^ knock-out mice are completely resistant to prion disease [12], knock-down of PrP^C^ in the brain using various strategies only delays the onset of clinical illness [13–15]. Thus, the discovery of alternate (non-PrP) targets for treating prion disease could be beneficial.

Despite extensive investigation, whether other proteins are required for prion replication to occur *in vivo* remains inconclusive [16]. Non-protein co-factors have been identified that enhance PrP^Sc^ production *in vitro*, but whether any of these molecules participate in prion replication *in vivo* remains unknown [17–22]. Evidence for the existence of non-PrP proteins involved in prion replication have been inferred from prion transmission studies [23]. Likewise, genome-wide association studies in CJD patients have linked several potential genes and their protein products to prion disease, including syntaxin-6, which appears to affect prion trafficking and secretion rather than playing a direct role in prion replication [24–26]. Moreover, while candidate prion replication modulators have been identified using genome-wide siRNA screens and by comparing the transcriptomes of prion-susceptible and prion-resistant cells, their broader relevance to prion disease remains to be established [27, 28].

Proteins that interact with PrP^C^ or reside in close spatial proximity to PrP^C^ within the cellular membrane have revealed clues about its molecular function [29–32]. Conceivably, a subset of these proteins may also influence prion replication. A frequently observed hit in PrP^C^ interactome studies are members of the protein disulfide isomerase (PDI) family. PDIs are thiol oxidoreductase chaperones that catalyze the oxidation, reduction, and isomerization of disulfide bonds between cysteine residues on client proteins during protein folding, which helps to maintain proteostasis [33–36]. In particular, the PDI family member prolyl 4-hydroxylase subunit beta (P4HB), also called PDIA1, resides in close spatial proximity to PrP^C^ at the cellular membrane in four different cell lines relevant to prion biology and in the brains of mice [29–31, 37]. Furthermore, P4HB and PrP^C^ can be co-immunoprecipitated in cultured cells, suggesting that the two proteins physically interact [38]. P4HB is organized into two active thioredoxin domains, *a* and *a*′, and two noncatalytic domains, *b* and *b′,* that are important for substrate recognition and binding [39–42]. In the active site of the *a* and *a*′ domains of P4HB, two cysteine residues are conserved within a Cys-Gly-His-Cys (CGHC) motif, and the *a* and *a*′ domains operate independently of each other [43]. Missense mutation or dysregulation of P4HB causes Cole-Carpenter syndrome as well as other illnesses including cancer, cardiovascular disease, and neurodegenerative diseases [44–52].

The disulfide bond in PrP^C^ plays a key role in stabilizing its structure and preventing it from misfolding and aggregating [53–56]. In cultured cell models, disulfide-free PrP mutants do not reach the cell surface and accumulate within the cytosol where they are largely insoluble [57–59]. Moreover, regulation of the disulfide bond in PrP^C^ is important for trafficking of PrP^C^ to the cell membrane and conversion of PrP^C^ to PrP^Sc^ at the cell surface [57–62]. Collectively, these findings highlight the importance of the disulfide bond for PrP biogenesis, implying that PDIs could play a role in prion pathogenesis. Two pan-PDI inhibitors increase levels of prion infection in cultured cells, implying that PDIs may be intrinsic inhibitors of prion replication [30]. Consistent with this notion, overexpression of PDIA3 reduces *de novo* prion infection in cultured cells and modestly improves survival in prion-infected mice [38, 63]. Moreover, P4HB upregulation has been reported in the brain of sporadic CJD patients and prion-infected hamsters [64–66]. However, whether P4HB regulates PrP^C^ and how it influences prion replication are not currently known.

In this study, we show that P4HB promotes formation of the disulfide bond in PrP. Stable knock-down of P4HB in cultured cells reduces PrP^C^ levels and dissuades prion infection. Furthermore, inhibition of the *a* domain in P4HB reduces PrP^C^ expression levels in uninfected cells and modulates PrP^Sc^ accumulation in prion-infected cells. These findings establish P4HB as a regulator of both PrP^C^ biogenesis and PrP^C^ to PrP^Sc^ conversion.

## Results

To investigate the role of P4HB in PrP^C^ homeostasis and prion replication, we utilized murine CAD5 cells. These cells are derived from the Cath.a.differentiated catecholaminergic cell line and are frequently used to study prion biology because they can be infected with many different strains of mouse prions [67, 68]. We first examined whether the disulfide bond in PrP^C^, which links cysteine residues 178 and 213 in mouse PrP (MoPrP), influences PrP^C^ biogenesis in CAD5 cells. For this purpose, we utilized CAD5-PrP^-/-^ cells, which permit examination of mutant PrP alleles without interference from endogenous wild-type (WT) MoPrP [69–71]. A disulfide-deficient MoPrP variant in which both cysteine residues are mutated to alanine (C178/213A) was generated (**Fig. 1a**). In transiently transfected CAD5-PrP^-/-^ cells, levels of disulfide-deficient MoPrP were significantly lower than WT MoPrP (**Fig. 1b, c**). Interestingly, when detecting PrP^C^ expression using the D18 antibody, which recognizes residues within the α-helical domain of PrP^C^ [72], significantly higher PrP signal was observed for WT MoPrP when samples were not reduced or boiled prior to immunoblotting. In contrast, sample reduction and boiling had only minimal effect on PrP signal for the C178/213A mutant. This stark signal difference for WT MoPrP was not observed when samples were probed with the antibody POM2, which recognizes the unstructured N-terminal domain of PrP^C^ [73] (**Fig. S1a**). This suggests that the absence of the disulfide bond in MoPrP prevents the α-helical domain from folding to its native state. In transfected CAD5-PrP^-/-^ cells, only a small amount of disulfide-deficient MoPrP reached the cell surface, as detected by immunofluorescent labeling of non-permeabilized cells (**Fig. 1d**). When cells were permeabilized prior to staining, higher signal for the C178/213A mutant was observed, suggesting that disulfide-deficient MoPrP accumulates within intracellular compartments. Whereas all three PrP^C^ glycoforms were observed when WT MoPrP was expressed, only a single PrP species was present for C178/213A-mutant MoPrP (**Fig. 1b**). To investigate whether this different banding pattern was caused by abnormal N-glycosylation, we treated lysates with PNGase F. Following removal of N-linked glycans, the molecular weights of full-length WT and C178/213A-mutant MoPrP were similar, indicating that N-glycosylation occurs abnormally when the PrP disulfide bond is absent (**Fig. 1e**). Interestingly, the C1 fragment, which results from physiological endoproteolytic processing of PrP^C^ in the vicinity of residues 110/111 [74], was not observed when the C178/213A mutant was expressed. Despite defective biosynthesis and trafficking, we did not observe an increase in the relevant amount of detergent-insoluble PrP species when C178/213A-mutant MoPrP was expressed (**Fig. S1b, c**). Collectively, these results show that absence of the disulfide bond in PrP alters PrP^C^ biogenesis in CAD5-PrP^-/-^ cells and are generally consistent with findings obtained using other cell lines [57–59].

**Figure 1.**
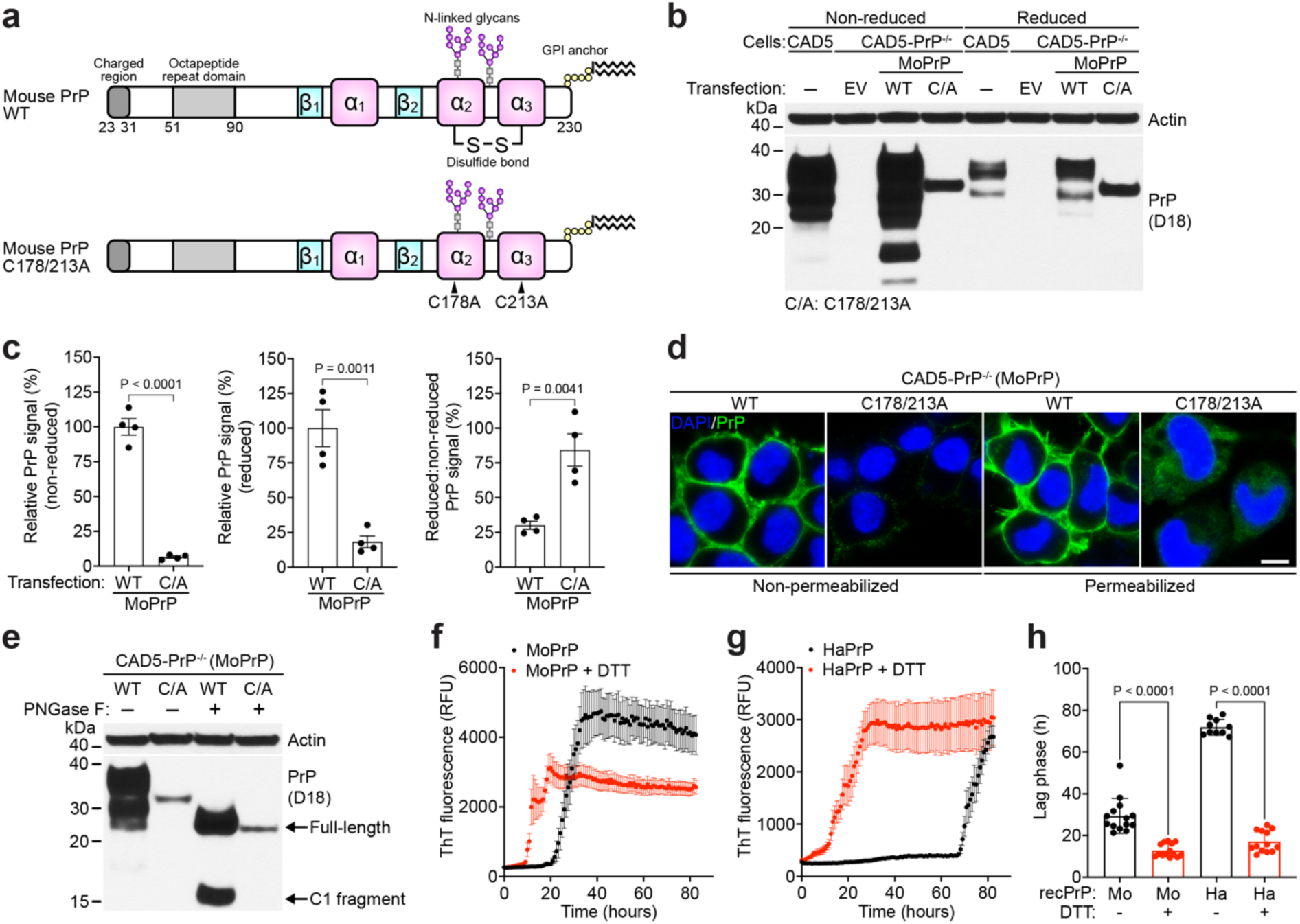
The disulfide bond in PrP^C^ increases protein stability and dissuades aggregation. (**a**) Schematic structures of WT mouse PrP (residues 23 to 230) and the disulfide-deficient mutant (C178/213A). (**b**) Representative immunoblots for PrP in cell lysates from CAD5-PrP^-/-^ cells transiently transfected with either empty vector (EV) or with WT or C178/213A-mutant MoPrP. Lysates from untransfected CAD5 cells are included as controls. In reduced conditions, lysates were treated with β-mercaptoethanol and boiled. The blot was probed with the anti-PrP antibody D18 and reprobed with an antibody against actin. (**c**) Quantification of PrP^C^ levels in non-reduced and reduced lysates from transiently transfected CAD5-PrP^-/-^ cells. Data are mean ± SEM from four independent samples. Statistical significance was assessed using unpaired two-tailed t tests. (**d**) Immunofluorescence images of CAD5-PrP^-/-^ cells transiently expressing either WT or C178/213A-mutant MoPrP. PrP^C^ expression (green) was revealed using the anti-PrP antibody POM1 and nuclei were stained with DAPI (blue). Scale bar = 10 μm (applies to all images). (**e**) Immunoblot for PrP in cell lysates from transfected CAD5-PrP^-/-^ cells with (+) or without (-) PNGase F treatment. The blot was probed with the anti-PrP antibody D18 and reprobed with an antibody against actin. (**f, g**) Representative aggregation curves for recombinant mouse PrP (MoPrP) (f) or hamster PrP (HaPrP) (g) when incubated alone (black; n = 14 for MoPrP, n = 10 for HaPrP) or in the presence of dithiothreitol (DTT) (red; n = 14 for MoPrP, n = 13 for HaPrP), as determined using a Thioflavin T (ThT) assay. Data are mean ± SEM. (**h**) Quantification of the lag phases for the ThT aggregation assays. Data are mean ± SEM. Statistical significance was assessed using one-way ANOVA followed by Tukey’s multiple comparisons test.

To examine whether loss of the disulfide bond in PrP^C^ promotes aggregation, we generated full-length recombinant (rec) MoPrP and hamster PrP (HaPrP) and then determined aggregation kinetics in the presence or absence of the reducing agent dithiothreitol using a Thioflavin T (ThT) assay performed under physiological conditions (i.e., at 37 °C in a buffer consisting of 10 mM sodium phosphate pH 7.3 and 135 mM sodium chloride). For both recMoPrP and recHaPrP, reduction of the PrP disulfide bond significantly accelerated the formation of ThT-positive PrP aggregates (**Fig. 1f-h**). Similar results have previously been obtained with recombinant human PrP [55, 56]. Thus, the disulfide bond stabilizes PrP^C^ and helps to prevent aggregation.

Having established the disulfide bond influences PrP stability and aggregation, we next asked whether P4HB modulates formation of the PrP disulfide bridge. For this purpose, we utilized recMoPrP and recP4HB in a PEG-maleimide (PEG-Mal) assay that labels free sulfhydryl groups in proteins (**Fig. 2a**). recMoPrP was first incubated with the reducing agent tris(2-carboxyethyl)phosphine (TCEP) at an elevated temperature to reduce the disulfide bond in PrP and cause it to unfold. Next, recMoPrP was incubated with or without recP4HB at 37 °C to allow re-folding and disulfide bond formation. If the two cysteine residues in recMoPrP remain reduced (i.e., the disulfide bond is not formed), PEG-Mal will covalently label recMoPrP, adding ∼10 kDa of molecular weight. If the two cysteine residues are oxidized (i.e., the disulfide bond has formed), recMoPrP will remain unlabeled. In the absence of recP4HB, the majority of recMoPrP was unlabeled but a signal for PEG-Mal-labeled recMoPrP was easily detectable, suggesting that the PrP disulfide bond reforms efficiently (**Fig. 2b**). When recP4HB was added, the amount of PEG-Mal-labeled recMoPrP was significantly decreased, implying that P4HB can assist with formation of the disulfide bond in PrP (**Fig. 2b, c**).

**Figure 2.**
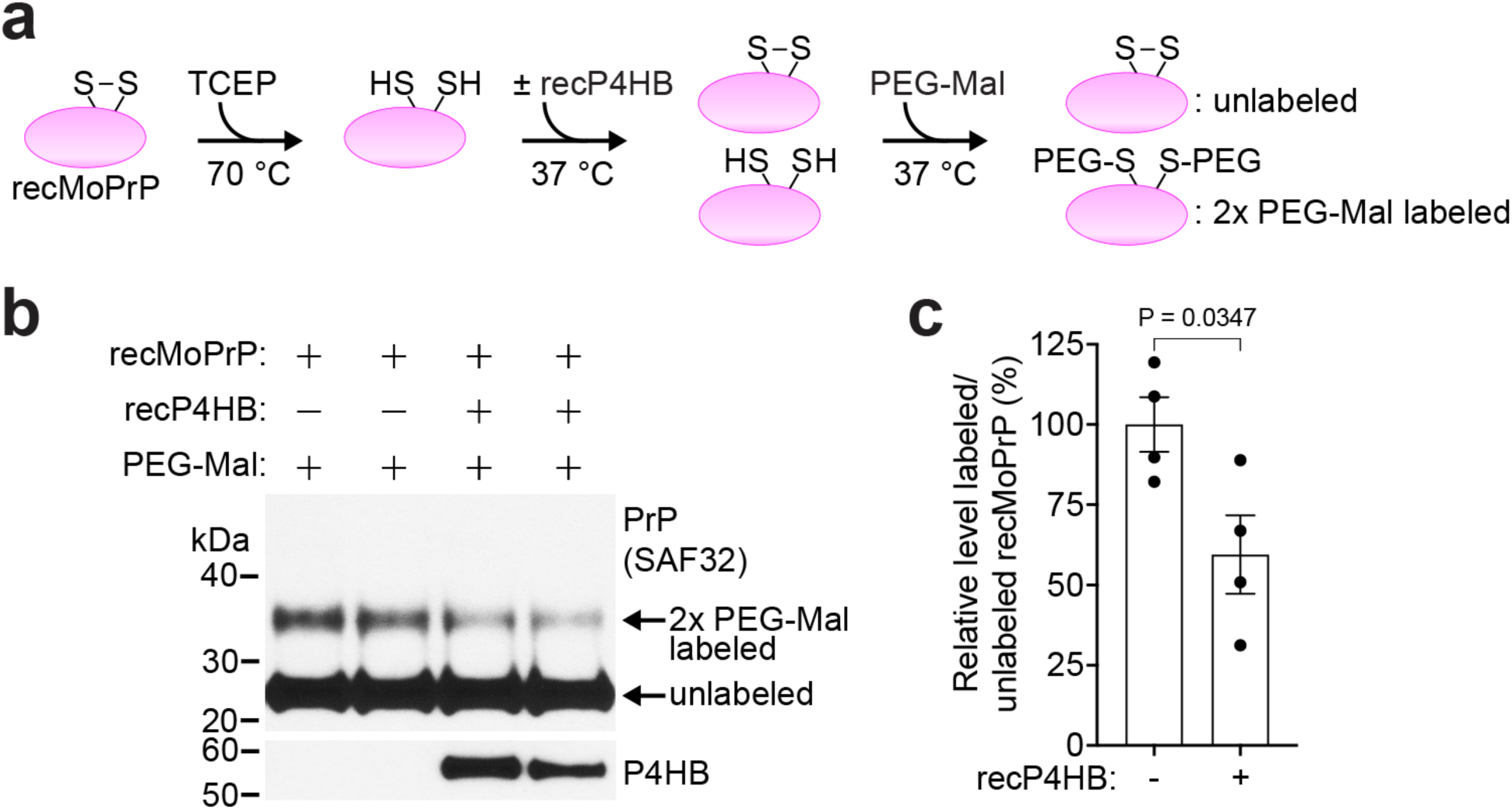
P4HB increases disulfide bond formation efficiency in PrP^C^. (**a**) Schematic of PEG-Maleimide (PEG-Mal) assay for assaying disulfide bond formation in recombinant mouse PrP (recMoPrP). (**b**) recMoPrP was reduced with tris(2-carboxyethyl)phosphine (TCEP), treated with (+) or without (-) recombinant P4HB (recP4HB), and then proteins were labeled with PEG-Mal. PEG-Mal labeled proteins were resolved by SDS-PAGE and analyzed by immunoblotting. The blot was probed with the anti-PrP antibody SAF32 or an anti-P4HB antibody. (**c**) Quantification of the relative ratio of PEG-Mal-labeled to unlabeled PrP for reactions performed in the absence (-) and presence (+) of recP4HB. Data are mean ± SEM from four independent replicates. Statistical significance was assessed using an unpaired, two-tailed t test.

Next, to determine the effect of P4HB depletion on PrP^C^ biogenesis, P4HB expression was reduced in CAD5 cells by transient transfection with siRNA, and then PrP^C^ expression was examined after three days. While transient knock-down of P4HB did not affect PrP^C^ levels, it did influence PrP glycosylation, with significantly higher levels of diglycosylated PrP and a concomitant reduction in mono- and unglycosylated PrP species in cells treated with P4HB siRNA (**Fig. 3a, b**). While ∼95% knock-down of P4HB was achieved, levels of the PDI family member PDIA3 were unaffected and levels of PDIA4 were only increased by ∼20% (**Fig. 3a, d**). Levels of GRP78 (also known as HSPA5 or BiP) were increased by ∼50% following P4HB knock-down, suggesting that transiently reducing P4HB levels causes ER stress (**Fig. 3d**) [75]. Transient knock-down of P4HB in CAD5 cells did not alter PrP^C^ trafficking nor did it increase levels of detergent-insoluble PrP species (**Fig. 3e-g**). However, levels of PrP in the conditioned medium increased following knock-down of P4HB, likely due to increased shedding of PrP from the cell membrane [74] (**Fig. S2a, b**).

**Figure 3.**
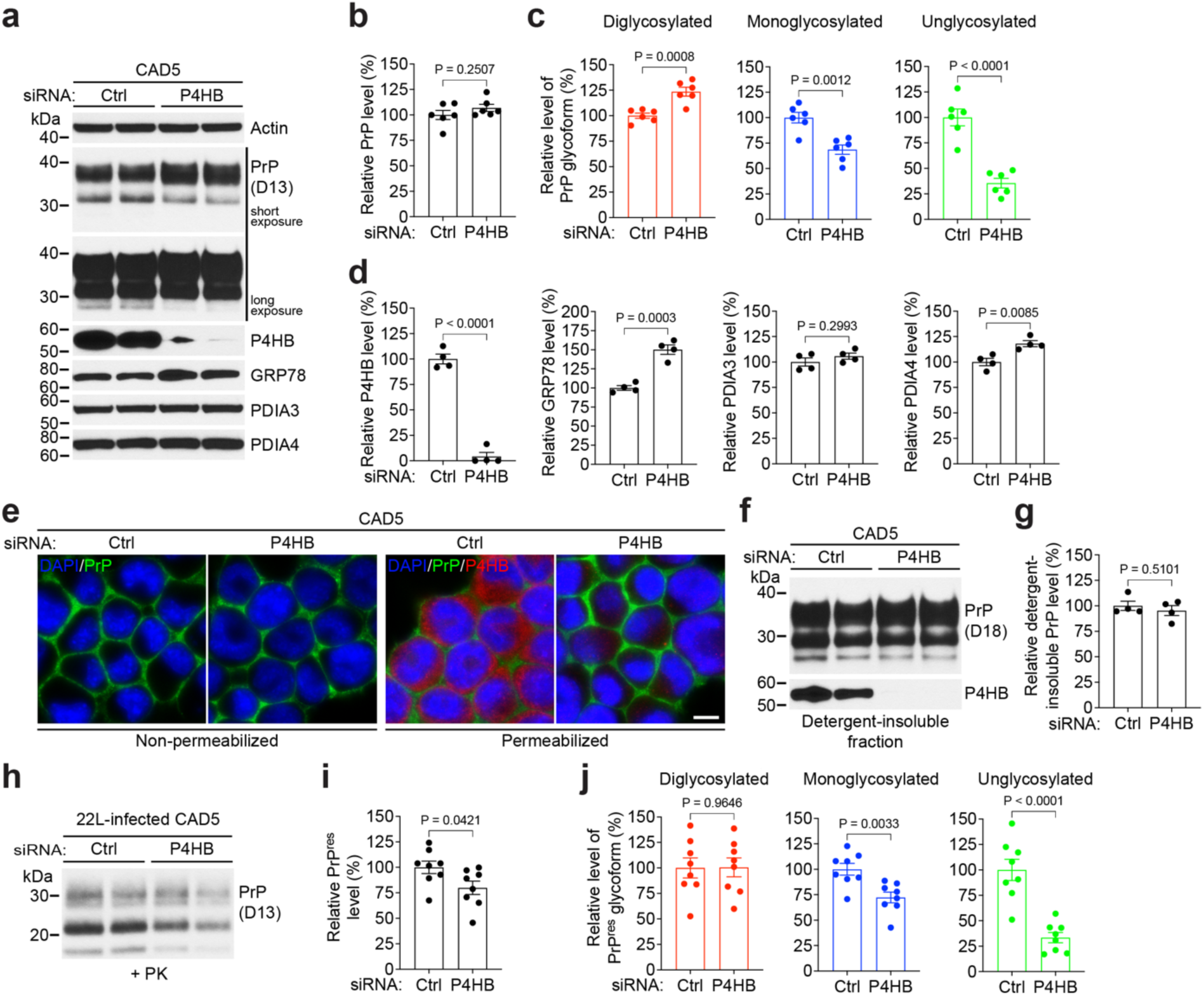
Transient knock-down of P4HB modulates PrP^C^ glycosylation and decreases PrP^res^ levels. (**a**) Representative immunoblots for PrP^C^, P4HB, GRP78, PDIA3 and PDIA4 in cell lysates from CAD5 cells transfected with control or P4HB siRNA for 72 h. (**b, c**) Quantification of PrP^C^ levels (b) and PrP^C^ glycoforms (c) in CAD5 cells transfected with control or P4HB siRNA (n = 6). (**d**) Quantification of P4HB, GRP78, PDIA3, and PDIA4 levels in CAD5 cells transfected with control or P4HB siRNA (n = 4). (**e**) Immunofluorescence images of CAD5 cells transfected with control or P4HB siRNA. PrP^C^ (green) and P4HB (red) were revealed using the anti-PrP antibody POM1 and an anti-P4HB antibody, and nuclei were stained with DAPI (blue). Scale bar = 10 μm (applies to all images). (**f**) Immunoblots of detergent-insoluble PrP and P4HB species in CAD5 cells transfected with control or P4HB siRNA for 72 h. (**g**) Quantification of detergent-insoluble PrP levels in CAD5 cells transfected with control or P4HB siRNA (n = 4). (**h**) Immunoblot of proteinase K (PK)-resistant PrP (PrP^res^) levels in 22L prion-infected CAD5 cells transfected with control or P4HB siRNA for 72 h. (**i, j**) Quantification of PrP^res^ levels (i) and PrP^res^ glycoforms (j) in 22L prion-infected CAD5 cells transfected with control or P4HB siRNA (n = 8). In panels b, c, d, g, i, and j, data are mean ± SEM and statistical significance was assessed using unpaired, two-tailed t tests.

To investigate if transient P4HB knock-down affects prion accumulation in CAD5 cells, we transfected CAD5 cells that had been stably infected with either the 22L or RML mouse prion strains with control or P4HB siRNA. After three days, levels of PK-resistant PrP (PrP^res^) in cell lysates were examined as a proxy for PrP^Sc^. In 22L prion-infected CAD5 cells, transient knock-down of P4HB significantly decreased PrP^res^ levels and altered its glycoform composition, with prominent decreases in the monoglycosylated and unglycosylated species (**Fig. 3h-j**). In RML prion-infected CAD5 cells, the monoglycosylated PrP^res^ species was also significantly decreased following transient P4HB knock-down whereas levels of total PrP^res^ were not as effected (**Fig. S3**). In summary, transient knock-down of P4HB in CAD5 cells modulates PrP^C^ glycosylation, which more strongly affects conversion of PrP^C^ to PrP^res^ in cells infected with 22L prions than in cells infected with RML prions.

We next investigated the effect of prolonged knock-down of P4HB on PrP^C^ biogenesis and prion infection. CAD5 cells were stably transfected with one of two distinct shRNAs targeting different regions in the P4HB gene as well as a control (non-targeting) shRNA. In the two P4HB knock-down lines, P4HB protein levels were decreased by ∼60-70% compared to the control line (**Fig. 4a, b**). Steady-state PrP^C^ levels were reduced by ∼30% when P4HB was stably knocked down. Unlike siRNA-mediated transient knock down of P4HB, stable knock down of P4HB did not elicit ER stress, as determined by a lack of GRP78 upregulation, nor did it cause increased production of PDIA3 or PDIA4 (**Fig. 4c**). In the P4HB knock-down lines, PrP^C^ was still predominantly present at the cell membrane (**Fig. 4d**). While absolute levels of detergent-insoluble PrP species were not increased in P4HB knock-down cells, when corrected for the decreased amount of total PrP, the relative proportion of insoluble PrP was slightly increased in one of the two knock-down lines (**Fig. 4e, f**). As upon transient knock-down of P4HB, shRNA-mediated stable knock-down of P4HB also caused an increased presence of secreted PrP species in the conditioned medium (**Fig. S2c, d**). To test whether prolonged P4HB knock down affects *de novo* prion infection, the stable knock-down CAD5 lines were challenged with RML or 22L prions. Levels of PrP^res^ were examined after five passages post-infection. In both P4HB knock-down lines, PrP^res^ levels were significantly decreased compared to the control line following infection with RML prions (**Fig. 4g, h**). In one of the two P4HB lines, PrP^res^ levels were significantly decreased following challenge with 22L prions (**Fig. 4i, j**). The extent of PrP^res^ reduction in P4HB knock down cells was greater for RML infections than 22L infections (∼35% vs. ∼17%). Therefore, unlike in the transient P4HB knock-down experiments where PrP^C^ glycosylation was altered, prolonged knock down of P4HB in CAD5 cells causes a decrease in PrP^C^ levels that hinders accumulation of the RML strain more so than the 22L strain following *de novo* prion infection.

**Figure 4.**
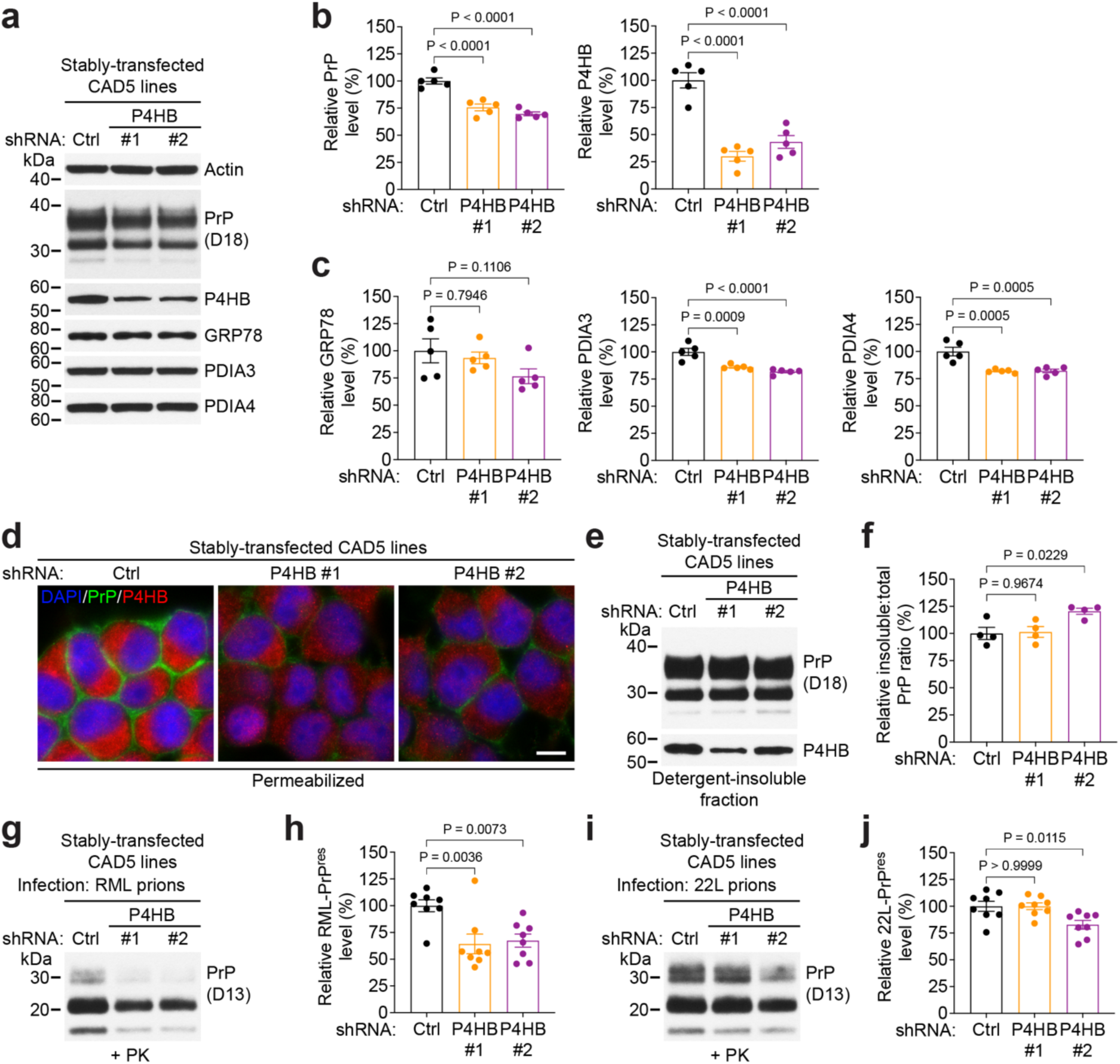
Stable knock-down of P4HB decreases PrP^C^ levels and hinders prion infection. (**a**) Representative immunoblots for PrP^C^, P4HB, GRP78, PDIA3 and PDIA4 in cell lysates from CAD5 cells stably transfected with control or one of two distinct P4HB shRNAs. (**b, c**) Quantification of PrP^C^ and P4HB levels (b) as well as GRP78, PDIA3, and PDIA4 levels (c) in CAD5 cells stably transfected with control or P4HB shRNAs (n = 5). (**d**) Immunofluorescence images of CAD5 cells stably transfected with control or P4HB shRNAs. PrP^C^ (green) and P4HB (red) were revealed using the anti-PrP antibody POM1 and an anti-P4HB antibody, and nuclei were stained with DAPI (blue). Scale bar = 10 μm (applies to all images). (**e**) Immunoblots of detergent-insoluble PrP and P4HB species in CAD5 cells stably transfected with control or P4HB shRNAs. (**f**) Quantification of the ratio of insoluble:total PrP in CAD5 cells transfected with control or P4HB shRNA (n = 4). (**g, i**) Immunoblots of PrP^res^ levels in CAD5 cells stably transfected with control or P4HB shRNAs and then infected with either RML (g) or 22L (i) prions. (**h, j**) Quantification of PrP^res^ levels in CAD5 cells stably transfected with control or P4HB shRNAs and then infected with either RML (h) or 22L (j) prions (n = 8). In panels b, c, f, h, and j, data are mean ± SEM and statistical significance was assessed using one-way ANOVA followed by Dunnett’s multiple comparisons test.

As an orthogonal approach to probing the role of P4HB in prion biosynthesis, we utilized a small molecule chemical inhibitor of P4HB. KSC-34 specifically targets Cys53 within the CGHC active site motif of the *a* domain within human P4HB and exhibits 30-fold selectivity for the *a* site (IC_50_ = 3.5 µM *in vitro*) compared to the *a*’ site (**Fig. 5a**) [76, 77]. To confirm that KSC-34 is also capable of selectively inhibiting the *a* domain of mouse P4HB, we treated CAD5 cells with 3.5 µM KSC-34 and then used a PEG-PCMal assay to analyze occupation of the P4HB active sites by the inhibitor. In the absence of KSC-34, the apparent molecular weight of P4HB increased by ∼20 kDa following PEG-PCMal treatment (**Fig. 5b**), suggesting that all four cysteine residues within the two active sites are reduced and accessible. However, when cells were first treated with KSC-34, the increase in apparent P4HB molecular weight following PEG-PCMal treatment was smaller, suggesting that binding of KSC-34 to one of the two P4HB active sites, presumably the *a* site, precludes labeling with PEG-PCMal. When similar experiments were performed for PDIA3 and PDIA4, no difference in PEG-PCMal labeling was observed upon treatment with KSC-34, suggesting that KSC-34 is specific for P4HB.

**Figure 5.**
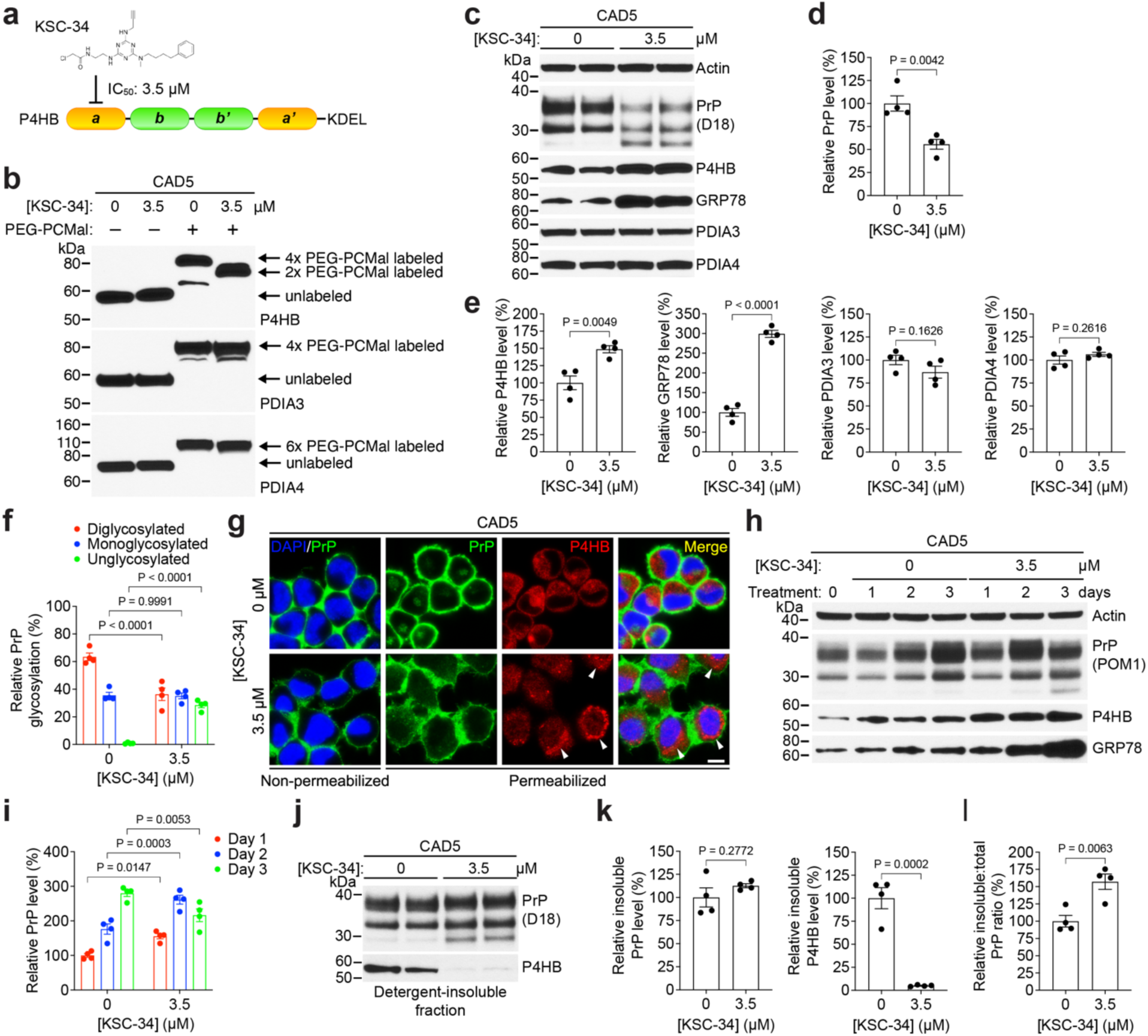
Inhibition of P4HB alters PrP^C^ glycosylation and decreases PrP^C^ levels in CAD5 cells. (**a**) Schematic structure of P4HB, which features catalytic *a* and *a′* domains (orange), non-catalytic ligand-binding domains *b* and *b′* (green), and an ER-retention motif (KDEL) at the C-terminus. The selective P4HB inhibitor, KSC-34, is 30-fold more selective for the *a* site than the *a′* site with an IC_50_ of ∼3.5 μM. (**b**) Lysates from CAD5 cells treated with 0 or 3.5 μM KSC-34 for 72 h were labeled with or without PEG-PCMal, and then PEG-PCMal labeled proteins were examined by immunoblotting. The blot was probed with antibodies to P4HB, PDIA3, and PDIA4. (**c**) Representative immunoblots for PrP^C^, P4HB, GRP78, PDIA3, and PDIA4 in cell lysates from CAD5 cells treated with 0 or 3.5 μM KSC-34 for 72 h. (**d, e**) Quantification of PrP^C^ (d) as well as P4HB, GRP78, PDIA3, and PDIA4 (e) levels in CAD5 cells treated with 0 or 3.5 µM KSC-34 for 72 h (n = 4). (**f**) Quantification of PrP^C^ glycoforms in CAD5 cells treated with 0 or 3.5 μM KSC-34 for 72 h (n = 4). (**g**) Immunofluorescence images of CAD5 cells treated with 0 or 3.5 µM KSC-34 for 72 h. The expression of PrP^C^ (green) and P4HB (red) were revealed using the anti-PrP antibody POM1 and an anti-P4HB antibody, and nuclei were stained with DAPI (blue). Scale bar = 10 μm (applies to all images). (**h**) Representative immunoblots of PrP^C^, P4HB, and GRP78 in cell lysates from CAD5 cells treated for 1, 2, or 3 days with 0 or 3.5 µM KSC-34. (**i**) Quantification of PrP^C^ levels in CAD5 cells treated for 1, 2, or 3 days with 0 or 3.5 µM KSC-34 (n = 4). (**j**) Representative immunoblots for detergent-insoluble PrP and P4HB species in cell lysates from CAD5 cells treated with 0 or 3.5 μM KSC-34 for 72 h. (**k**) Quantification of detergent-insoluble PrP and P4HB levels in CAD5 cells treated with 0 or 3.5 μM KSC-34 for 72 h (n = 4). (**l**) Quantification of the ratio of insoluble:total PrP in CAD5 cells treated with 0 or 3.5 µM KSC-34 for 72 h (n = 4). All data are mean ± SEM. In panels d, e, k, and l, statistical significance was assessed using unpaired two-tailed t tests. In panels f and i, statistical significance was assessed using a two-way ANOVA followed by Šídák’s multiple comparison test.

CAD5 cells were treated with KSC-34 for 3 days and then levels of PrP^C^ and P4HB in cell lysates were examined by immunoblotting (**Fig. 5c**). PrP^C^ levels were reduced by ∼45% upon treatment of CAD5 cells with KSC-34, and P4HB levels were increased by ∼50% (**Fig. 5c-e**). This decrease in PrP^C^ levels was observed in both reduced and non-reduced samples as well as when PrP^C^ was detected using an antibody that recognizes the unstructured PrP N-terminal domain (**Fig. S4**). Levels of GRP78 were strongly increased following KSC-34 treatment, indicating that KSC-34 elicits ER stress, whereas levels of the other PDI family members PDIA3 and PDIA4 remained unchanged (**Fig. 5e**). In addition to decreased PrP^C^ levels, the glycosylation of PrP was also altered in the presence of KSC-34 (**Fig. 5c, f**). Levels of unglycosylated PrP were significantly increased at the expense of the diglycosylated PrP glycoform, which was significantly decreased (**Fig. 5f**). Immunofluorescence experiments on CAD5 cells treated with KSC-34 revealed that the majority of PrP was still present at the cell membrane (**Fig. 5g**). However, treatment of CAD5 cells with KSC-34 caused a punctate intracellular distribution of P4HB. Following treatment of CAD5 cells for one or two days, KSC-34 initially caused an increase in PrP^C^ levels, followed by a decrease in PrP^C^ levels and an increase in unglycosylated PrP on day three (**Fig. 5h, i**). PrP^C^ levels were also decreased in non-dividing CAD5 cells that had been differentiated to a more neuron-like state by serum deprivation (**Fig. S5**). Unlike in P4HB knock-down cells, treatment with KSC-34 caused a significant decrease in secreted PrP levels (**Fig. S6a, b**). This may potentially be explained by a decrease in the precursor and mature forms of the ADAM10 protease (**Fig. S6c, d**), which is responsible for PrP shedding from the cell membrane [78, 79]. Despite a decrease in total PrP^C^ levels, the relative proportion of detergent-insoluble PrP species was significantly increased in KSC-34-treated CAD5 cells (**Fig. 5j-l**). Interestingly, despite total P4HB levels also increasing following KSC-34 treatment, levels of insoluble P4HB were significantly decreased (**Fig. 5j, k**). Collectively, these results suggest that inhibition of the *a* domain in P4HB perturbs PrP homeostasis in CAD5 cells.

Next, we tested the effect of partial P4HB inhibition on pre-existing PrP^Sc^ levels in prion-infected CAD5 cells. Prion infection, either with the RML or 22L strains, had no effect on P4HB levels (**Fig. S7**). Unexpectedly, given that KSC-34 decreased PrP^C^ levels in uninfected CAD5 cells, treatment of either RML- or 22L-prion infected CAD5 cells with KSC-34 for three days caused a significant increase in both total PrP and PrP^res^ levels (**Fig. 6a-c**). PrP^res^ levels were increased by ∼50% and ∼25% in RML- and 22L-prion infected cells, respectively. As in uninfected cells, KSC-34 treatment caused an increase in P4HB levels in prion-infected CAD5 cells (**Fig. 6d**). The effect of KSC-34 on PrP^res^ levels was dose-dependent in RML-infected CAD5 cells (**Fig. 6e, f**). Continuous treatment of RML-infected CAD5 cells for four passages resulted in an ∼3-fold increase in PrP^res^ levels (**Fig. 6g, h**). Thus, partial inhibition of P4HB in CAD5 cells potentiates prion infection.

**Figure 6.**
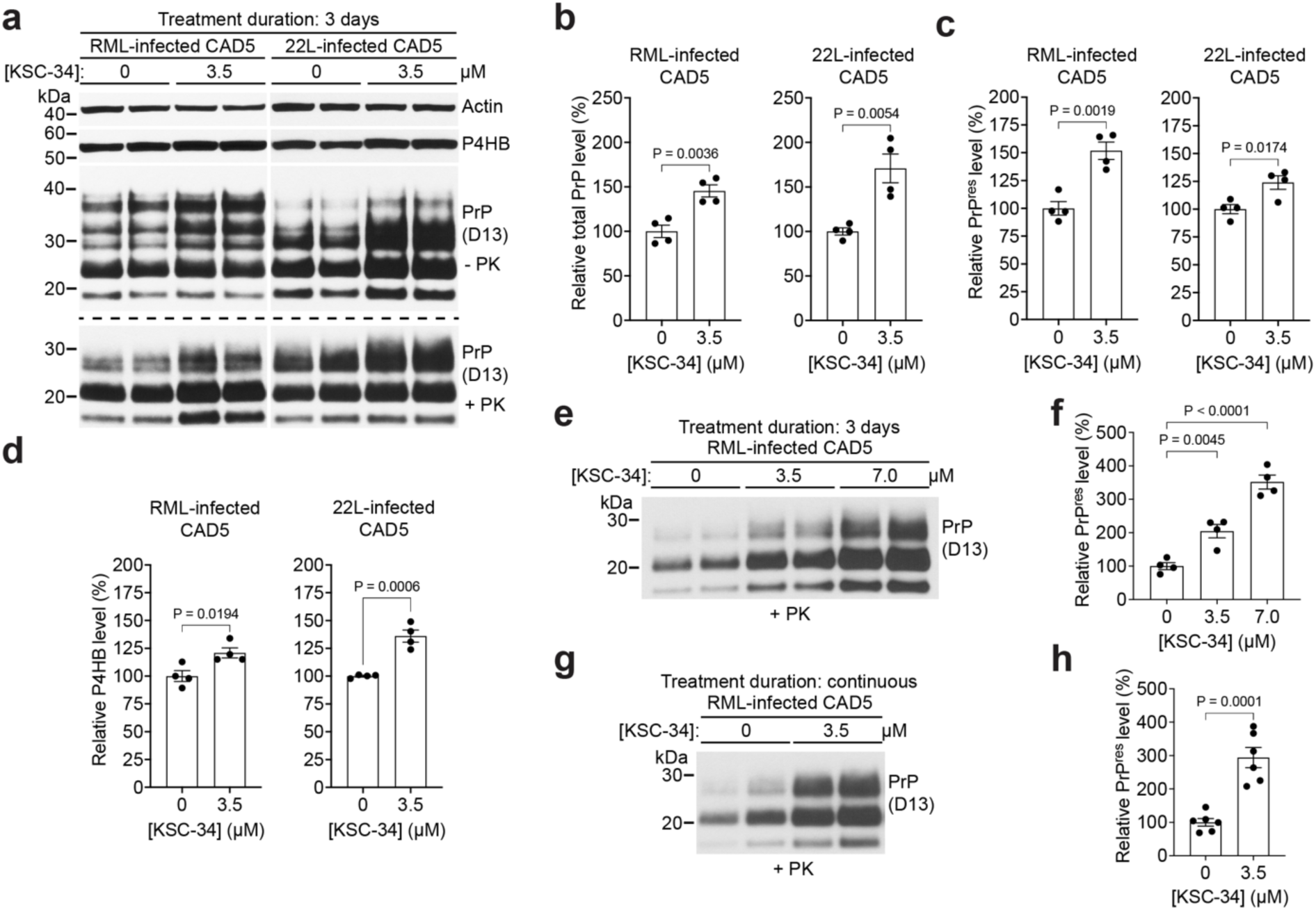
Inhibition of P4HB increases PrP^res^ levels in prion-infected CAD5 cells. (**a**) Representative immunoblots for total PrP, P4HB and proteinase K (PK)-resistant PrP^Sc^ (PrP^res^) levels in lysates from either RML or 22L prion-infected CAD5 cells treated with 0 or 3.5 μM KSC-34 for 72 h. The blots were probed with the anti-PrP antibody D13, an anti-P4HB antibody, and an anti-actin antibody. (**b-d**) Quantification of total PrP (b), PrP^res^ (c), and P4HB (d) levels in RML or 22L prion-infected CAD5 cells treated with 0 or 3.5 μM KSC-34 (n = 4 independent replicates). (**e**) Representative immunoblot of PrP^res^ levels in RML prion-infected CAD5 cells treated with 0, 3.5, or 7 µM KSC-34 for 72 h. (**f**) Quantification of PrP^res^ levels in RML prion-infected CAD5 cells treated with 0, or 3.5, or 7 µM KSC-34 (n = 4 independent replicates). (**g**) Representative immunoblot of PrP^res^ levels in RML prion-infected CAD5 cells cultured in the continuous presence of either 0 or 3.5 µM KSC-34 for 4 passages. (**h**) Quantification of PrP^res^ levels in RML prion-infected CAD5 cells treated with 0 or 3.5 µM KSC-34 for X days (n = 6 independent replicates). In panels b-d, f, and h, graphs display mean ± SEM. Statistical significance in panels b-d, and h was assessed using unpaired, two-tailed t tests. In panel f, statistical significance was assessed using one-way ANOVA followed by Dunnett’s multiple comparisons test.

Conversion of PrP^C^ to PrP^Sc^ is believed to occur on the cell surface or within the endolysosomal pathway [80, 81]. Despite the presence of a C-terminal KDEL ER-retention sequence in P4HB, a proportion of P4HB can reach the cell surface [82, 83]. In cerebrospinal fluid, secreted P4HB levels correlate with total and phosphorylated tau levels and are increased in Alzheimer’s disease patients [84]. Furthermore, it has been shown that cell-surface P4HB is detectable in N2a neuroblastoma cells [30]. To determine whether a proportion of P4HB is also present on the cell surface of CAD5 cells, we performed a cell surface protein biotinylation assay. Following labeling of cell surface proteins using a membrane-impermeant biotinylation reagent and pull-down of biotin-labeled protein using streptavidin, signal for both PrP^C^ and P4HB was observed in the eluate fraction, indicating that a fraction of both proteins is localized on the cell surface (**Fig. 7a**). The absence of actin signal in the eluate fraction indicates that biotin labeling of P4HB did not arise from labeling of intracellular proteins due to compromised membrane integrity. To examine the effect of cell-surface P4HB on PrP^Sc^, we generated a secreted P4HB variant lacking the C-terminal ER retention motif (ΔKDEL) (**Fig. 7b**). We also generated versions of WT and ΔKDEL P4HB in which the catalytic cysteine residues in the *a* site are mutated to serine (C55/58S). In CAD5 cells transiently transfected with each of the four P4HB variants, levels of P4HB in cell lysates were significantly increased (**Fig. 7c, d**). However, levels of PrP^C^ were unchanged. In conditioned medium, P4HB was detectable in cells transfected with the ΔKDEL P4HB variants but not in cells transfected with WT P4HB (**Fig. 7e**). Transient transfection of both RML- and 22L-prion infected CAD5 cells with the ΔKDEL variant resulted in an increase in PrP^res^ levels (**Fig. 7f, g**). For RML prions, the increase was ∼50% and for 22L prions, the increase was ∼35%. For both prion strains, an increase in PrP^res^ was also observed for the ΔKDEL variant in which the *a* domain catalytic site was inactivated, potentially pointing to an important role for the *a*’ domain catalytic site. While the *a* and *a*’ CGHC active sites in P4HB exhibit approximately equal catalytic activity, they are functionally non-equivalent [43, 85]. In summary, an increase in secreted but not ER-resident P4HB causes an increase in prion replication in CAD5 cells.

**Figure 7.**
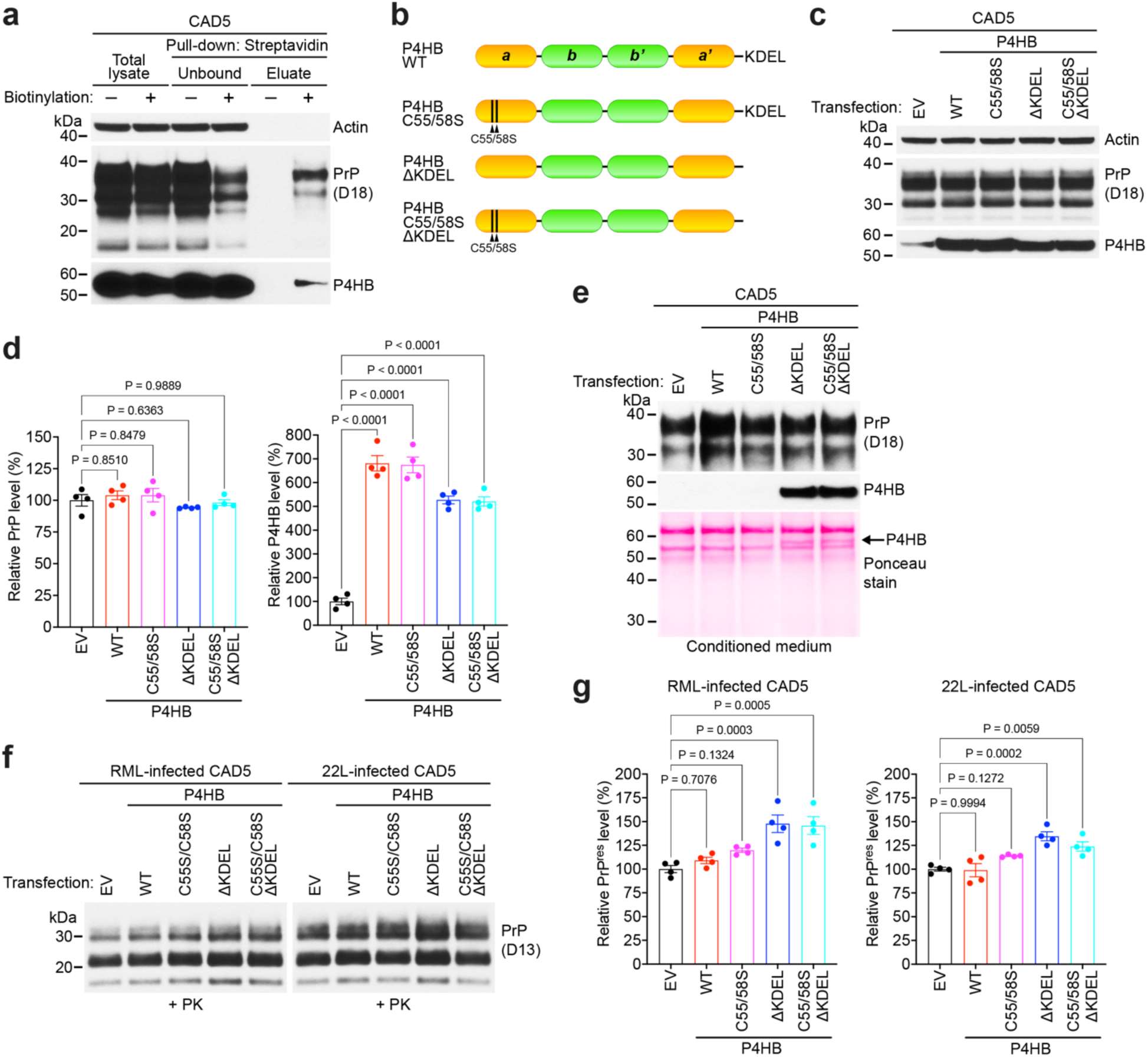
Transient transfection with a secreted version of P4HB increases PrP^res^ levels in CAD5 cells. (**a**) Immunoblots for PrP^C^ and P4HB in total cell lysates as well as unbound and eluate fractions following incubation with streptavidin agarose from CAD5 cells subjected to cell surface biotinylation (+) or mock (-) treatment. (**b**) Schematic structures of WT mouse P4HB as well as mutants with a catalytically inactive *a* site (C55/58S) and/or deletion of the C-terminal ER retention signal (ΔKDEL). (**c**) Immunoblot of PrP^C^, P4HB, and actin levels in cell lysates from CAD5 cells transiently transfected with empty vector (EV) or with WT or mutant P4HB for 48 h. (**d**) Quantification of PrP^C^ and P4HB levels in lysates from transiently transfected CAD5 cells (n = 4 independent replicates). (**e**) Immunoblot of secreted PrP and P4HB levels in the conditioned medium from CAD5 cells transiently transfected with EV or with WT or mutant P4HB for 48 h. As a control, the membrane was stained for total protein following transfer using Ponceau S. (**f**) Representative immunoblots for proteinase K (PK)-resistant PrP^Sc^ (PrP^res^) levels in lysates from either RML or 22L prion-infected CAD5 cells transiently transfected with EV or with WT or mutant P4HB for 48 h. (**g**) Quantification of PrP^res^ levels in transiently transfected RML or 22L prion-infected CAD5 cells (n = 4 independent replicates). In panels d and g, graphs display mean ± SEM and statistical significance was assessed using one-way ANOVA followed by Dunnett’s multiple comparisons test.

Finally, since the ΔKDEL variant must transit through the secretory pathway, where it could conceivably influence PrP^C^ and/or PrP^Sc^ biogenesis before it reaches the cell surface, we wanted to test whether application of P4HB directly to the cell surface can influence prion replication. Therefore, we transiently transfected CAD5-PrP^-/-^cells with the two ΔKDEL P4HB variants and then collected the conditioned medium. CAD5-PrP^-/-^ cells were used for these experiments to prevent any confounding influence from secreted forms of PrP. As expected, secreted P4HB was only observed in the conditioned medium from CAD5-PrP^-/-^ cells transfected with the ΔKDEL P4HB variants (**Fig. 8a**). The conditioned medium from the transiently transfected CAD5-PrP^-/-^ cells was then applied to 22L prion-infected CAD5 cells (**Fig. 8b**). Levels of PrP^res^ were modestly but significantly increased following application of conditioned medium from cells expressing ΔKDEL P4HB (**Fig. 8c, d**). In this case, the result was specific to the ΔKDEL P4HB variant with an intact catalytic site within the *a* domain. Therefore, cell-surface P4HB can have a positive influence on prion infection. However, we did not observe a similar effect in RML prion-infected CAD5 cells (**Fig. S8**).

**Figure 8.**
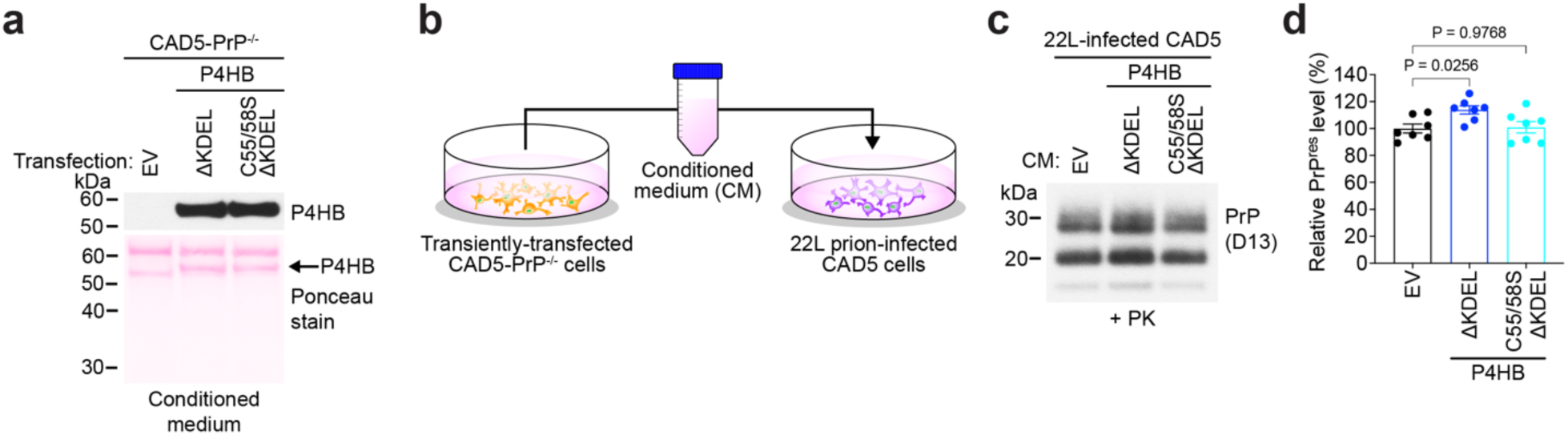
Conditioned medium containing P4HB increases PrP^res^ levels in 22L prion-infected CAD5 cells. (**a**) Immunoblot of secreted P4HB levels in the conditioned medium (CM) from CAD5-PrP^-/-^ cells transiently transfected with empty vector (EV) or with mutant P4HB variants lacking the C-terminal ER retention signal. As a control, the membrane was stained for total protein following transfer using Ponceau S. (**b**) Schematic of the experiment in which CM from transiently transfected CAD5-PrP^-/-^ cells is applied to 22L prion-infected CAD5 cells for 72 h. (**c**) Representative immunoblot of proteinase K (PK)-resistant PrP^Sc^ (PrP^res^) levels in lysates from 22L prion-infected CAD5 cells treated with CM from CAD5-PrP^-/-^ cells transiently transfected with EV or secreted P4HB variants. (**d**) Quantification of PrP^res^ levels in lysates from 22L prion-infected CAD5 cells treated with CM from CAD5-PrP^-/-^ cells transiently transfected with EV or secreted P4HB variants (n = 7 independent replicates). The graph displays mean ± SEM and statistical significance was assessed using one-way ANOVA followed by Dunnett’s multiple comparisons test.

## Discussion

The identification of non-PrP proteins that influence the PrP^C^ to PrP^Sc^ transition has been a long-sought goal of the field, as they may provide clues towards the molecular mechanism of prion replication and could constitute novel therapeutic targets for treating prion diseases. However, the discovery of such proteins has been challenging, with no definitive prion modulators emerging to date, although high-throughput screens are now revealing new candidates [27, 86]. In previous studies, the protein disulfide isomerase P4HB/PDIA1 has consistently been identified as a protein that either interacts directly with PrP^C^ or resides in close spatial proximity to PrP^C^ within the cell [30, 31, 37, 38]. Given that P4HB is known to play an important role in protein folding and proteostasis within the ER, either by assisting with formation and rearrangement of disulfide bonds or by acting as molecular chaperone, a putative role for P4HB in the biogenesis of PrP^C^ and/or PrP^Sc^ seemed plausible. Here, we have demonstrated that perturbation of P4HB influences both PrP^C^ levels and prion replication in cultured cells. We propose that P4HB is required for efficient prion replication *in vivo* and plays a complex role in PrP homeostasis by 1) helping to maintain PrP^C^ stability and steady-state PrP^C^ levels; and 2) acting as a chaperone that directly augments the refolding of PrP^C^ into PrP^Sc^ at the cell surface (**Figure 9**).

**Figure 9.**
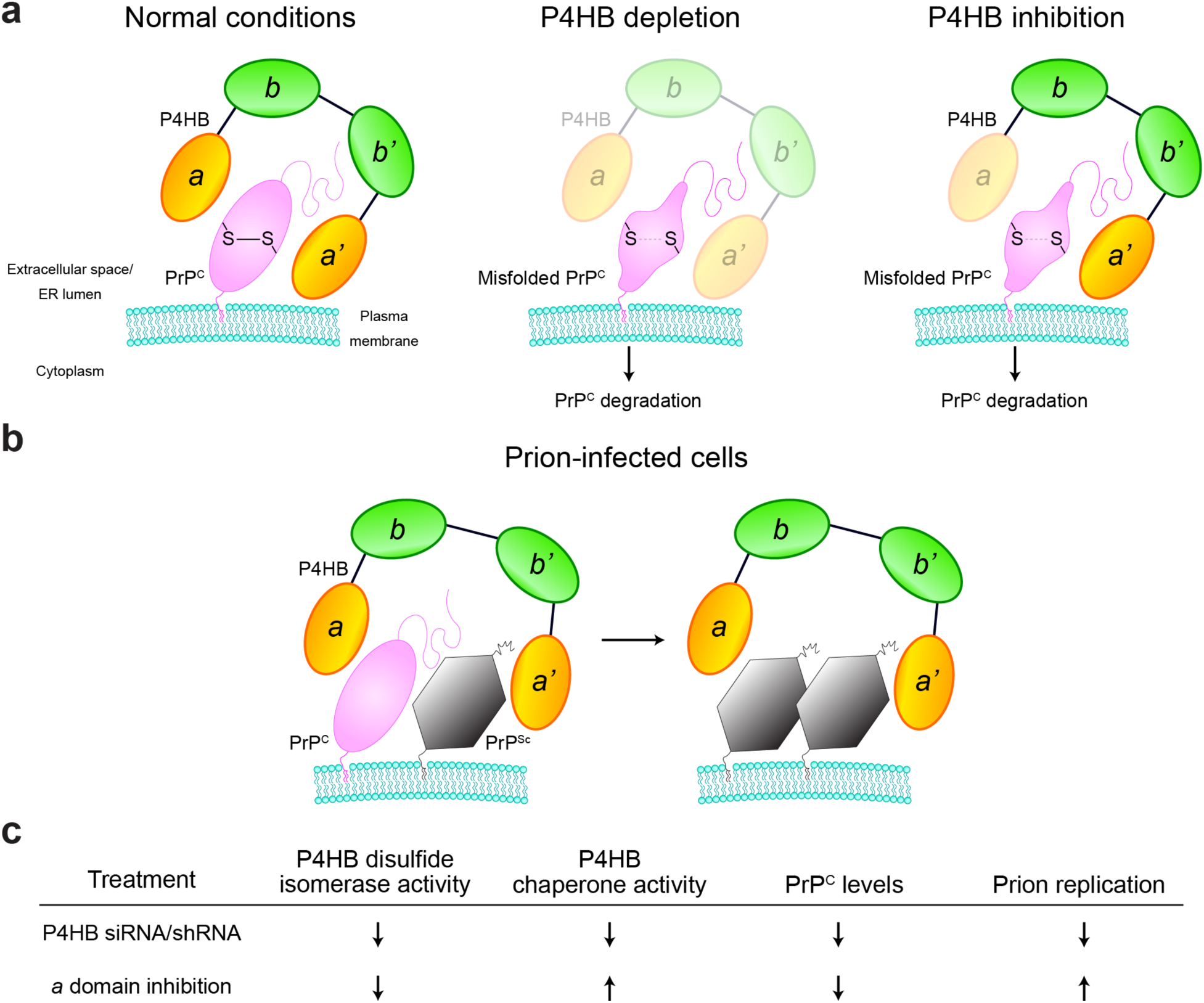
Model outlining a potential dual role for P4HB in modulating PrP^C^ levels and prion replication. (**a**) P4HB helps to maintain the disulfide bond in PrP^C^. When P4HB is depleted or its disulfide isomerase activity is inhibited, the disulfide bond in PrP^C^ forms less efficiently causing PrP^C^ to become misfolded and degraded, which leads to lower steady-state PrP^C^ levels. (**b**) P4HB functions as a chaperone that potentiates prion replication. In prion-infected cells, P4HB may act as a scaffold that facilitates interactions between PrP^C^ and PrP^Sc^. (**c**) When P4HB levels are decreased, both the isomerase and chaperone activities of P4HB are decreased, leading to lower PrP^C^ levels and less prion replication. When the catalytic *a* domain is inhibited, the chaperone activity of P4HB increases, leading to increased prion replication despite lower PrP^C^ levels.

CAD5 cells in which P4HB levels were knocked down by stable expression of shRNAs exhibited lower PrP^C^ levels and accumulated less PrP^Sc^ following prion infection. In other experiments, a disulfide-free PrP variant accumulated to lower levels in cells, which is suggestive of decreased protein stability, and P4HB assisted with formation of the PrP disulfide bond *in vitro*. A simple explanation for these observations is that P4HB helps to maintain PrP^C^ stability by ensuring proper formation and maintenance of the disulfide bond. When P4HB levels are decreased, PrP^C^ misfolds more often or becomes less stable leading to increased turnover and lower steady-state PrP^C^ levels as well as a concomitant decrease in PrP^Sc^ levels following prion infection (**Figure 9a**). However, two observations suggest that P4HB can also directly influence PrP^Sc^ and prion replication, independent of its effect on PrP^C^. First, application of conditioned medium containing extracellular P4HB to 22L prion-infected CAD5 cells or expression of a secreted P4HB mutant in RML or 22L prion-infected CAD5 cells increased PrP^Sc^ levels. Second, in several assays we observed that perturbation of P4HB levels had strain-specific effects on PrP^Sc^. Most notably, treatment with conditioned medium containing extracellular P4HB modulated PrP^res^ levels in cells infected with 22L, but not RML prions. This is consistent with the finding that overexpression of PDIA3 increases PrP^C^ levels yet affects levels of 22L, but not RML prions in cultured cells [63].

P4HB is known to function within the ER as both a “foldase” that assists with protein folding via its disulfide isomerase activity and a chaperone “holdase” that helps to prevent protein misfolding, activities that are regulated by phosphorylation of P4HB at Ser357 [87]. During prion replication, perhaps extracellular P4HB acts as a scaffold that optimally positions PrP^C^ for templated conversion into PrP^Sc^ by prion aggregates (**Figure 9b**). Lowering the disulfide isomerase and chaperone activities of P4HB simultaneously using siRNA/shRNA would decrease conversion of PrP^C^ to PrP^Sc^ by both lowering PrP^C^ levels and reducing template-directed misfolding (**Figure 9c**). In contrast, lowering the disulfide isomerase activity of P4HB alone via pharmacological inhibition or mutation of the *a* domain may unlock P4HB’s chaperone activity, allowing it to act as a better scaffold for mediating prion replication. Indeed, inhibition of the P4HB *a* domain using KSC-34 significantly increased PrP^res^ levels following treatment of prion-infected cells, as did overexpression of a secreted version of P4HB with a catalytically inactive *a* domain. It should be noted that, because PrP^C^ levels initially increase following KSC-34 treatment, the increase in PrP^res^ could also partially result from a temporary uptick in prion replication due to an increase in PrP^C^ substrate. Alternatively, prolonged inhibition of the P4HB *a* domain may strongly destabilize PrP^C^, potentially causing partial unfolding and creating a PrP species that can more readily convert into PrP^Sc^ in prion-infected cells. The precise molecular details of how P4HB’s individual domains and its dual molecular activities contribute to regulating PrP^C^ and PrP^Sc^ biogenesis will require further investigation.

While P4HB likely exerts a direct effect on PrP^C^ and/or PrP^Sc^, other indirect mechanisms could also be involved. Induction of ER stress in cultured cells results in an increase in both insoluble PrP species as well as PrP^res^ [88]. We found that transient knock-down of P4HB or partial inhibition of P4HB using KSC-34 induced ER stress in CAD5 cells, as judged by an increase in GRP78 levels. However, stable knock-down of P4HB in CAD5 cells decreased PrP^C^ levels and reduced the efficiency of *de novo* prion infection in the absence of an increase in GRP78 levels. Thus, induction of ER stress due to reduced P4HB activity is unlikely to be the major driver of the observed effects on PrP biosynthesis. We also found that levels of secreted PrP were modulated by P4HB knock-down and inhibition. Secreted forms of PrP^C^ have been postulated to function as endogenous inhibitors of prion replication, possibly by trapping PrP^Sc^ in the extracellular space [89]. Indeed, we found that transient or stable knock-down of P4HB in CAD5 cells increases production of secreted PrP, and this correlated with a decrease in PrP^res^ levels in prion-infected cells. Conversely, levels of secreted PrP were decreased following P4HB inhibition, and this correlated with an increase in PrP^res^ levels in prion-infected CAD5 cells. Therefore, the observed effects on PrP^Sc^ in CAD5 cells when P4HB activity is reduced may be at least partially mediated by a change in extracellular PrP levels.

A limitation of this study is that the P4HB knockdown/inhibition experiments were only performed using an immortalized cultured cell line. In mice, P4HB has been successfully knocked out in several cell types, including myeloid cells, platelets, β-cells, and osteoblasts [90–94], whereas whole-body knockout of P4HB has been reported to be embryonic lethal [94]. Post-natal, whole-body knockout of P4HB can be achieved, but these mice exhibit severe defects in spermatogenesis, leading to sterility in male animals [95]. To the best of our knowledge, the consequences of either full or partial knockout of P4HB in the brain or specific brain cell types has not yet been investigated. Going forward, it will be important to generate mice lacking P4HB in the brain to better understand the role of P4HB in PrP^C^ biogenesis and prion replication within an intact animal. Another limitation is that although there are more than 20 protein disulfide isomerase family members in humans [36], we only focused on P4HB in this study. Although P4HB is the most commonly identified PDI family member in PrP interactome studies, other PDIs such as PDIA3, PDIA4, and PDIA6 have also been found to reside in close spatial proximity to PrP in cells [30]. Thus, it is conceivable that other PDIs, perhaps acting in synergy with P4HB, may also influence prion disease-relevant biology, as has been demonstrated for PDIA3 [63]. However, we did not notice major compensatory changes in the levels of PDIA3 or PDIA4 following knockdown or inhibition of P4HB in CAD5 cells, which is consistent with what has been observed in mice [95]. Therefore, it should be feasible to individually evaluate the contribution of each PDI family member to prion disease pathogenesis.

The development of therapeutics for treating or curing animal and human prion diseases remains an urgent priority. Anti-prion small molecules can extend survival following infection of mice with mouse prion strains but have so far remained ineffective at treating human prions or preventing spontaneous prion formation [96–102]. Down-regulation of PrP^C^ levels has emerged as the dominant therapeutic strategy, as it is predicted to be effective against all prion strains, including those from humans. Indeed, antisense oligonucleotides that decrease PrP^C^ levels are currently in clinical trial, and several other strategies that diminish PrP^C^ levels via gene editing or transcriptional modulation are in various stages of pre-clinical development [13–15, 103–105]. However, complete reduction of PrP^C^ in the brain using these strategies is likely impossible, suggesting that it may be necessary to combine other therapeutic strategies with PrP^C^ reduction. While no fully validated non-PrP therapeutic targets currently exist for prion disease, drugs targeting various cellular processes or proteins have shown some limited efficacy in prion-infected mice [106–108]. Whether knock-down of P4HB may have therapeutic benefit in prion disease when used alone or in combination with PrP-targeting strategies remains to be determined. Given that P4HB is important for maintaining proteostasis and likely influences the folding and trafficking of many proteins that transit through the secretory pathway, it is likely that targeting P4HB may generate unwanted side effects, at least under certain conditions. Thus, it may be necessary to fine-tune the extent of P4HB reduction to not disrupt critical cellular pathways. Alternatively, small molecules that specifically disrupt interactions between PrP and P4HB may reduce PrP^C^ levels and/or prion conversion without directly affecting P4HB’s normal activities.

## Methods

### Materials and chemicals

The following antibodies were used in this study: anti-PrP antibodies SAF32 (Cayman Chemical #189720), POM1 (Sigma #MABN2285), and POM2 (Sigma #MABN2298) as well as the recombinant humanized anti-PrP Fabs HuM-D18 and HuM-D13 (respectively referred to as D18 and D13 for simplicity) [72]; anti-P4HB (Abcam #ab137110), anti-PDIA3 (ThermoFisher #PA585083), anti-PDIA4 (ThermoFisher #PA530321), Anti-GRP78 (ThermoFisher #MA5-35606), anti-ADAM10 (Abcam #ab124695), and anti-Actin (Sigma #A5060). D13 was a generous gift from Stanley Prusiner. Horseradish peroxidase-conjugated secondary antibodies were purchased from Bio-Rad (#172-1011, #172-1019) and ThermoFisher (#31414). KSC-34 was obtained from MedChemExpress (#HY-117570).

### Plasmid constructs

Expression plasmids encoding mouse PrP were generated by cloning the respective *Prnp* open reading frames between the BamHI and Xbal sites of the vector pcDNA3 [109]. The C178/213A mutant based on mouse PrP was generated by site-direct mutagenesis. To prepare the P4HB expression plasmids, WT mouse P4HB cDNA (#MG50638-UT, SinoBiological) or the ΔKDEL mutant was inserted between the BamHI and NotI sites of pcDNA3. The C55/58S mutants were generated by site-direct mutagenesis. All constructs were confirmed by DNA sequencing.

### Cell culture

Murine CAD5 cells, which were obtained from Charles Weissmann, are a subclone of the catecholaminergic CAD line [67, 68]. CAD5 cells were cultured in growth medium [Opti-MEM medium (Thermo Fisher #31985088) containing 10% (v/v) fetal bovine serum and 2 mM GlutaMAX (Gibco #35050061)]. To passage the cells, they were first washed with PBS (ThermoFisher #14190144) and then treated with enzyme-free cell dissociation reagent (Millipore Sigma #S-014-B). The cells were incubated for 1-3 minutes at 37 °C, dissociated, and then plated in fresh growth medium. To prepare differentiated CAD5 cells, CAD5 cells were cultured in DMEM/F-12 medium (Invitrogen #11330032) with 2 mM GlutaMAX. The medium was changed every two days over a period of five days. CAD5-PrP^-/-^ cells, which were generated by CRISPR/Cas9 gene editing, were cultured as described previously [69]. All cell lines were cultured at 37°C in a 5% CO_2_ atmosphere at constant humidity.

### Prion strains

The mouse prion strains RML and 22L were derived from the brains of terminally ill prion-infected non-transgenic C57BL/6 mice. To generate prion-infected cell lines, CAD5 cells were incubated with 0.2% brain homogenate which contains either RML or 22L for 72 h. After the incubation, the cells were passaged several times, and infection status was analyzed by detection of PK-resistant PrP. To generate prion-infected cellular homogenates, RML or 22L prion-infected cells were cultured in 10-cm tissue culture plates. After reaching confluence, the cells were washed in cold PBS and then scraped into a small volume of PBS and collected into CK14 soft tissue homogenization tubes (Bertin Technologies #P000912-LYSK0-A) supplemented with additional 0.5 mm zirconia beads (BioSpec #11079105Z). The cells were homogenized using the Minilys apparatus (Bertin Technologies) using 3 cycles of 60 s at maximum speed, with a 10 min incubation on ice between each cycle. Following homogenization, cellular homogenates were aliquoted and stored at −80 °C.

### Transient transfections

Uninfected CAD5 or CAD5-PrP^-/-^ cells were plated at 3.0 x 10^5^ cells/well in a 6-well plate. The following day, 2.5 µg of plasmid DNA was mixed with 5 µL of Lipofectamine 2000 (ThermoFisher #11668019) in 200 µL of serum-free Opti-MEM medium, and the mixture was added to the cells and incubated for 24 h, after which the medium was replaced with fresh growth medium. After 48 h post-transfection, the cells were lysed for analysis of protein expression. Prion-infected CAD5 cells were seeded at 1.0 x 10^6^ cells in 6-cm dishes. The following day, 6.0 µg of plasmid DNA was mixed with 12 µL of Lipofectamine 2000 in 300 µL of serum-free Opti-MEM medium, and the mixture was added to the cells and incubated for 24 h. After 24 h from the transfection, the medium was replaced with fresh growth medium, and the cells were lysed for analysis of PK-resistant PrP 48 h post-transfection. For siRNA transfections, uninfected or prion-infected CAD5 cells were plated at 4.0 x 10^5^ cells in 6-cm dishes. The following day, 50 pmol of control or P4HB siRNA (QIAGEN #1027280, #1027417[Mm_P4hb_1 FlexiTube siRNA]) was mixed with 15 µL of Lipofectamine RNAiMAX (ThermoFisher #13778075) in 300 µL of serum-free Opti-MEM medium, and the mixture was added to the cells and incubated for 24 h. After 24 h post-transfection, the medium was replaced with fresh growth medium, and the cells were lysed for analysis of proteinase K-resistant PrP 72 h post-transfection.

### Generation of stably transfected cell lines

For generation of CAD5 cells stably transfected with P4HB shRNA, CAD5 cells were seeded at a density 5.0 x 10^5^cells/well in a 6-well plate. The following day, 2 µg of control or P4HB shRNA (ORIGENE #TR501569) was mixed with 4 µL of Lipofectamine 2000 in 200 µL of serum-free Opti-MEM medium, and the mixture was added to the cells and incubated for 24 h. After 24 h post-transfection, the cells were washed with PBS and detached using the enzyme-free dissociation reagent. The transfected cells were then transferred into a 6-cm dish and cultured in medium containing 5 µg/mL puromycin for 5 days, and then stably transfected cells were expanded. No clonal selection was performed. To ensure stable knock-down of P4HB, CAD5 cells which were stably transfected with P4HB shRNA were maintained in growth medium containing 1 µg/mL puromycin.

### Cellular prion infections

For *de novo* prion infection experiments, stably transfected CAD5 cells were plated in 24-well plates at a density of 5.0 x 10^4^ cells/well. After 24 h from seeding the cells, they were cultured in 500 µL of growth media with 100 µg of cellular homogenate containing the RML or 22L mouse prion strains for 72 h. After 72 h, the cells were washed with PBS and then passaged in 12-well plates. Cells were continuously passaged in 12-well plates for at least 3 passages, and then scaled up to 6-cm dishes prior to lysis for the analysis of infection status and PK-resistant PrP. As negative controls, cells were exposed to cellular homogenate from uninfected cells.

### Cell lysis and immunoblotting

Cultured cells were washed with DPBS and then lysed using lysis buffer, which consists of 50 mM Tris-HCl pH 7.4, 150 mM NaCl, 0.5% (w/v) sodium deoxycholate, 0.5% (v/v) NP-40 and protease inhibitor cocktail (Sigma #11873580001). The cell lysate was incubated on ice for 30 min, and then centrifuged at 5,000x *g* for 10 min to remove the insoluble cell debris. Protein concentrations in the supernatant were quantified using the BCA assay (ThermoFisher #23227), and then samples were diluted as required into 1x Bolt LDS sample buffer (ThermoFisher #B0007). Unless otherwise indicated, all immunoblot samples were prepared under non-reduced conditions in the absence of β-mercaptoethanol. For reduced conditions, 2.5% v/v) β-mercaptoethanol was added and samples were boiled at 95 °C for 10 min. Samples were electrophoresed on Bolt Bis-Tris gels (ThermoFisher #NW00100BOX and #NW04120BOX) using the MES buffer system. Following SDS-PAGE, proteins were transferred to Immobilon-P PVDF membranes (Millipore #IPVH00010), which were then blocked with blocking buffer [5 % (w/v) skim milk in Tris-buffered saline containing 0.05 % (v/v) Tween-20 (TBST)]. Membranes were then incubated overnight at 4 °C with antibodies diluted in blocking buffer. The following day, membranes were washed 3 times with TBST (5 min per wash), and then incubated with horseradish peroxidase-conjugated secondary antibodies diluted in blocking buffer for 1 h at room temperature. The membranes were again washed 6 times with TBST (10 min per wash). The blots were developed using Western Lightning ECL pro (Revvity #NEL122001EA) and then chemiluminescent signal was captured by exposure to HyBlot CL x-ray film (Thomas Scientific #1141J52). Immunoblotting data were quantified using NIH ImageJ software.

### Detergent insolubility assays

Cultured cells were lysed with lysis buffer containing protease inhibitor cocktail. For analysis of detergent-insoluble PrP and P4HB species, 100 µg of cell lysate was adjusted to a final volume of 100 µL of lysis buffer. The samples were subjected to ultracentrifugation at 100,000× *g* for 1 h at 4 °C. Following ultracentrifugation, the supernatant was removed by gentle aspiration, and the pellet was resuspended in 20 µL of 1x LDS sample buffer. The samples were boiled for 10 min at 95 °C, and then 100 µg of detergent-insoluble protein was loaded onto Bolt Bis-Tris gels and analyzed by immunoblotting.

### Proteinase K digestions

Cultured cells were lysed with lysis buffer that lacks the protease inhibitor cocktail. For analysis of PrP^res^, 500 µg of cell lysate was adjusted to a final volume of 200 µL of lysis buffer containing 50 µg/mL proteinase K (PK) (ThermoFisher #EO0491) for a final PK-to-protein ratio of 1:50. Samples were digested for 1 h at 37 °C with shaking at 600 rpm, and the digestions were then terminated by the addition of PMSF to a concentration of 2 mM. Sarkosyl was added to a final concentration of 2% (v/v), and the samples were then subjected to ultracentrifugation at 100,000× *g* for 1 h at 4 °C. Following ultracentrifugation, the supernatant was removed by gentle aspiration, and the pellet was resuspended in 1x LDS sample buffer. The samples were boiled for 10 min at 95 °C and then stored at −80 °C. 100 µg of PK-digested protein was loaded onto Bolt Bis-Tris gels and analyzed by immunoblotting.

### PNGase F digestions

20 μg of cell lysate from CAD5-PrP^-/-^ cells transiently transfected with either WT or C178/213A-mutant MoPrP was mixed with 1 μL of Glycoprotein Denaturing Buffer (10x) and dH_2_O in a total reaction volume of 10 μL. The mixture was denatured for 10 min at 95 °C, and then samples were mixed with 2 μL of GlycoBuffer 2 (10x), 2 μL of 10% NP-40, 6 μL of dH_2_O and 1 μL of PNGase F (New England BioLabs #P0708S). The mixture was incubated for 1 h at 37 °C, and then the samples were diluted as required into 1x Bolt LDS sample buffer. Deglycosylated samples were analyzed by immunoblotting as described above.

### KSC-34 treatment

KSC-34 was dissolved in DMSO. Uninfected CAD5 cells were plated at a density of 2.0-4.0 x 10^5^ cells in 6-cm dishes. Prion-infected CAD5 cells were seeded at a density of 4.0-8.0 x 10^5^ cells in 6-cm dishes. The following day, cells were treated with 0, 3.5, or 7.0 µM KSC-34 for 24-72 h and then the cells were lysed for analysis of protein expression or PK-resistant PrP. In the experiments where prion infection occurred under continuous KSC-34 treatment, uninfected CAD5 cells were plated in 24-well dishes at a density of 5.0 x 10^4^ cells/well. The following day, the cells were treated with 0 or 3.5 µM KSC-34 in the presence of 25 µg of cellular homogenate containing RML mouse prions. After 72 h of incubation, the cells were washed with PBS and then passaged into 24-well dishes. The cells were then passaged three times in 24-well plates depending on cell confluency. At the fourth passage, the cells were scaled up to 6-cm dishes and cultured in the presence of 0 or 3.5 µM KSC-34 until they were collected for the analysis of *de novo* prion accumulation. The medium containing KSC-34 was changed every three days.

### Immunofluorescence microscopy

CAD5 cells or CAD5-PrP^-/-^ cells were plated in 24-well dishes with a #1.5 glass-like polymer coverslip bottom (Cellvis #P24-1.5P) coated with poly-D-lysine (ThermoFisher #A3890401). Prior to the addition of cells, the coated wells were thoroughly washed twice with PBS. For the KSC-34 treatment experiments, CAD5 cells were plated at density of 8.0 x 10^4^ cells/well. The following day, cells were treated with 0 or 3.5 µM KSC-34 for 72 h. For the siRNA transfection experiments, CAD5 cells were seeded at density of 5.0 x 10^4^ cells/well. The following day, the cells were transfected with 5 pmol of siRNA. After 24 h post-transfection, the medium was replaced with fresh growth medium, and then cells were cultured for 48 h. Stable cell lines were plated at a density of 1.0 x 10^5^ cells/well and were cultured for 48 h until they reached 80–90% confluency. Cells were fixed with 4% (v/v) paraformaldehyde for 15 min and then washed twice with PBS. Permeabilization was performed with 0.1% Triton-X (diluted in PBS) for 10 min at 4 °C. After washing with PBS, the cells were blocked in 3% bovine serum albumin (diluted in PBS) for 1 h at room temperature. Cells were then incubated with the anti-PrP antibody POM1 (1:500 dilution) and the anti-P4HB antibody (0.8 µg/mL) overnight at 4°C in PBS containing 3% bovine serum albumin. The following day, the cells were washed twice with PBS and then incubated with Alexa Fluor conjugated secondary antibodies (ThermoFisher #A-11029, #A-11001, 1:500 dilution in PBS containing 3% bovine serum albumin) for 2 h at room temperature. Subsequently, cells were washed 2 additional times with PBS. After the final wash, the cells were incubated with DAPI (1 µg/mL in PBS containing 3% bovine serum albumin) for 10 min and then washed with PBS. The cells were stored in PBS and then imaged using a Zeiss LSM880 confocal microscope.

### Acetone concentration of secreted proteins

In KSC-34 treated CAD5 cells, after 48 h post-KSC-34 treatment, the cells were washed with PBS, and then the cells were cultured in serum-free Opti-MEM containing 0 or 3.5 µM KSC-34 for 24 h. In the transient P4HB overexpression experiments, the transfected CAD5 cells were washed with PBS after 24 h post-transfection, and then the cells were cultured in serum-free Opti-MEM for 24 h. In the siRNA-mediated transient P4HB knockdown experiments, the transfected CAD5 cells were washed with PBS after 48 h post-transfection, and then the cells were cultured in serum-free Opti-MEM for 24 h. The stable cell lines were cultured for 48 h until they reached 70-80% confluency, washed with PBS, and then the cells were cultured in serum-free Opti-MEM for 24 h. The conditioned medium was collected and then centrifuged at 1,000x *g* for 10 min to remove cell debris. The supernatant was stored at −80 °C. To precipitate proteins, 300 µL of conditioned medium was mixed with 1.2 mL of cold acetone, and the mixture was incubated overnight at −30 °C. The mixture was centrifuged at 16,000x *g* for 10 min at 4 °C, and then the pellet was dried for 30 min on ice to completely remove the acetone, and then lysis buffer was added to each sample. The proteins were eluted from the pellet for 30 min with shaking at 1,500 rpm, and then the samples were centrifuged at 16,000x *g* for 10 min at 4 °C. Protein concentrations in the supernatant were quantified using the BCA assay, and the samples were prepared for immunoblotting. PVDF membranes with transferred proteins ware stained with Ponceau S solution (Bioshop #PON002.500) for 15 min, and then membranes were washed twice with dH_2_O for 2.5 min for destaining. After washing with TBST, the secreted proteins in conditioned medium were analyzed by immunoblotting as described above.

### Conditioned medium treatment in prion-infected cells

CAD5-PrP^-/-^ cells were seeded at 3.0 x 10^6^ cells in 10-cm dishes. The following day, 16 µg of plasmid DNA was mixed with 32 µL of Lipofectamine 2000 in 1 mL of serum-free Opti-MEM medium, and the mixture was added to the cells and incubated for 24 h. After 24 h from the transfection, the medium was replaced with 17 mL of fresh growth medium, and then the medium was collected after 48 h from transfection. The collected medium was centrifuged at 1,000x *g* for 10 min to remove cell debris, and then 16.5 mL of the supernatant was used as the conditioned medium and was stored at 4 °C. Prion-infected CAD5 cells were seeded at 1.6 x 10^6^ cells in 6-cm dishes. The following day, the medium was replaced with 5 mL of conditioned medium, and then the prion-infected cells were cultured for 3 days. The conditioned medium was changed every 24 h. Cells were then lysed for analysis of PK-resistant PrP.

### Biotinylation assay

CAD5 cells were plated in a 15-cm dish and were cultured for 48 h until they reached 80–90% confluency. The cells were washed with 20 mL of PBS and were incubated with 10 mL of PBS containing Sulfo-NHS-SS-Biotin (ThermoFisher #44390) for 10 min at room temperature according to the manufacturer’s instructions. Biotin treated cells were washed twice with 10 mL of ice-cold TBS and collected into a 50 mL centrifuge tube. The samples were centrifuged at 500x *g* for 5 min at 4 °C, and then the supernatant was discarded. The cells were lysed with 600 μL of lysis buffer containing protease inhibitor cocktail, and then the cell lysate was incubated on ice for 30 min and was centrifuged at 15,000x *g* for 10 min at 4 °C to remove the insoluble cell debris. The lysate was divided into 100 μL as the total cell lysate and 500 μL for the isolation of biotin-labeled proteins. The biotin-labeled cell surface proteins were isolated with NeutrAvidin Agarose according to the manufacturer’s instructions (ThermoFisher #44390), and then the unbound-proteins were collected. The bound proteins were eluted with 250 μL of elution buffer containing 10 mM DTT. The samples were diluted as required into 1x Bolt LDS sample buffer. Biotin-labeled samples were analyzed by immunoblotting as described above.

### Production of recombinant PrP and ThT assays

Untagged, full-length recombinant mouse PrP (residues 23-230) or hamster PrP (residues 23-231) were expressed in *E. coli* Rosetta2 (DE3) cells as then purified as previously described [71]. Briefly, inclusion bodies were solubilized using 8 M guanidine hydrochloride and then recombinant PrP was captured using Ni-NTA Superflow agarose beads (Qiagen #30430). PrP was refolded on-column using a 4-hour gradient from 6 to 0 M guanidine hydrochloride and then eluted using a gradient of 0 to 500 mM imidazole. Purified recombinant PrP was dialyzed into 10 mM sodium phosphate pH 5.8, filtered, and then stored at −80 °C. For the ThT assays, recombinant PrP was dialyzed into 10 mM sodium phosphate pH 7.3 buffer and then ultracentrifuged at 100,000× *g* for 1 h at 4 °C immediately prior to use to remove any pre-existing aggregates. The reaction mixtures contained 0.1 mg/mL recombinant PrP, 135 mM NaCl, and 10 µM ThT in a final volume of 100 µL of 10 mM sodium phosphate pH 7.3 in black, clear-bottom 96-well microplates (ThermoFisher #265301). In some reactions, 10 mM DTT was added. The assays were performed using a BMG CLARIOstar microplate reader set at 37 °C with cycles of 4 min shaking and 1 min rest/read. ThT fluorescence was read every 5 min using the following settings: excitation: 444 ± 5 nm; emission: 485 ± 5 nm; gain: 1600. Aggregation curves were fit using a variable slope (four parameter) sigmoidal model in GraphPad Prism (version 10.6.0) and then lag phases were calculated using the equation T_50_ – (1/2k) where T_50_ is the time at which 50% of maximal fluorescence is reached and k is the Hill slope.

### Gel shift assay

Purified recombinant mouse PrP and His-tagged recombinant mouse P4HB (Sino Biological #50638-M08H) were dialyzed into 50 mM Tris-HCl pH 7.4, 150 mM NaCl. 0.1 µg of recombinant PrP was reduced using 2.5 mM TCEP and 0.1% SDS for 10 min at 70 °C. Reduced recombinant PrP was then treated with 0.1 µg of recombinant P4HB for 1 h at 37 °C in the presence of TCEP and SDS. Following treatment with recombinant P4HB, the mixture was incubated with 1.0 mM PEG-Mal (Sigma #63187) for 30 min at 37 °C, and then samples were diluted as required into 1x Bolt LDS sample buffer as described above. The cell lysate was incubated with 1.0 mM PEG-PCMal (Dojindo Molecular Tech. Inc. #SB20-01) for 30 min at 37 °C, and then samples were diluted as required into 1x Bolt LDS sample buffer. The samples were loaded onto the Bolt Bis-Tris gels. Following SDS-PAGE, the gels were treated with UV radiation at 365 nm for 20 min before proteins were transferred to PVDF membranes. Proteins labelled with PEG-Mal or PEG-PCMal were analyzed by immunoblotting as described above.

### Statistical analysis

All data was assumed to be normally distributed. When comparing the means of two samples, two-tailed paired or unpaired t tests were used. When comparing the means of three or more samples, one-way ANOVA followed by either Dunnett’s (when comparing means to a control sample) or Tukey’s (when comparing all sample means to each other) multiple comparisons test was used. For some experiments, samples were compared using two-way ANOVA followed by Šídák’s multiple comparison test. All statistical analysis was performed using GraphPad Prism software (version 10.6.0) with a significance threshold of *P* < 0.05.

## Declarations

### Availability of data and material

All data generated or analyzed during this study are included in this published article.

### Author’s contributions

The study was conceived and designed by GA, GSU, and JCW. GA performed most of the experiments and conducted most of the data analysis. The ThT aggregation assays in Figure 1f-h were performed by HA and ZP. The first draft of the manuscript was written by GA and JCW, and all authors commented on previous versions of the manuscript. All authors read and approved the final manuscript.

---

**Figure S1.**
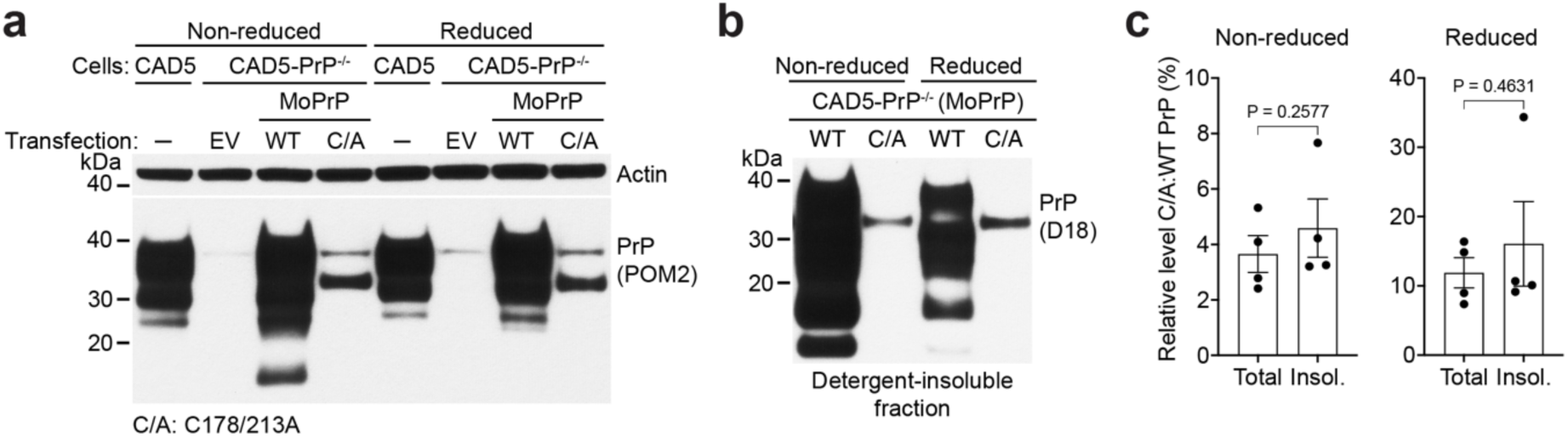
Absence of the disulfide bond does not increase levels of detergent-insoluble PrP. (**a**) Representative immunoblots for PrP in cell lysates from CAD5-PrP^-/-^ cells transiently transfected with either empty vector (EV) or with WT or C178/213A-mutant MoPrP. Lysates from untransfected CAD5 cells are included as controls. In reduced conditions, lysates were treated with β-mercaptoethanol and boiled. The blot was probed with the anti-PrP antibody POM2 and reprobed with an antibody against actin. (**b**) Representative immunoblot of detergent-insoluble PrP species in cell lysates from CAD5-PrP^-/-^ cells transiently transfected with either WT or C178/213A-mutant MoPrP. Samples were either reduced or left unreduced prior to loading. (**c**) Quantification of the ratio of C178/213A:WT PrP signal for non-reduced and reduced samples in total cell lysate and detergent-insoluble (insol.) fractions from CAD5-PrP^-/-^cells transiently transfected with either WT or C178/213A-mutant MoPrP (n = 4 independent replicates). The graph displays mean ± SEM and statistical significance was assessed using paired two-tailed t tests.

**Figure S2.**
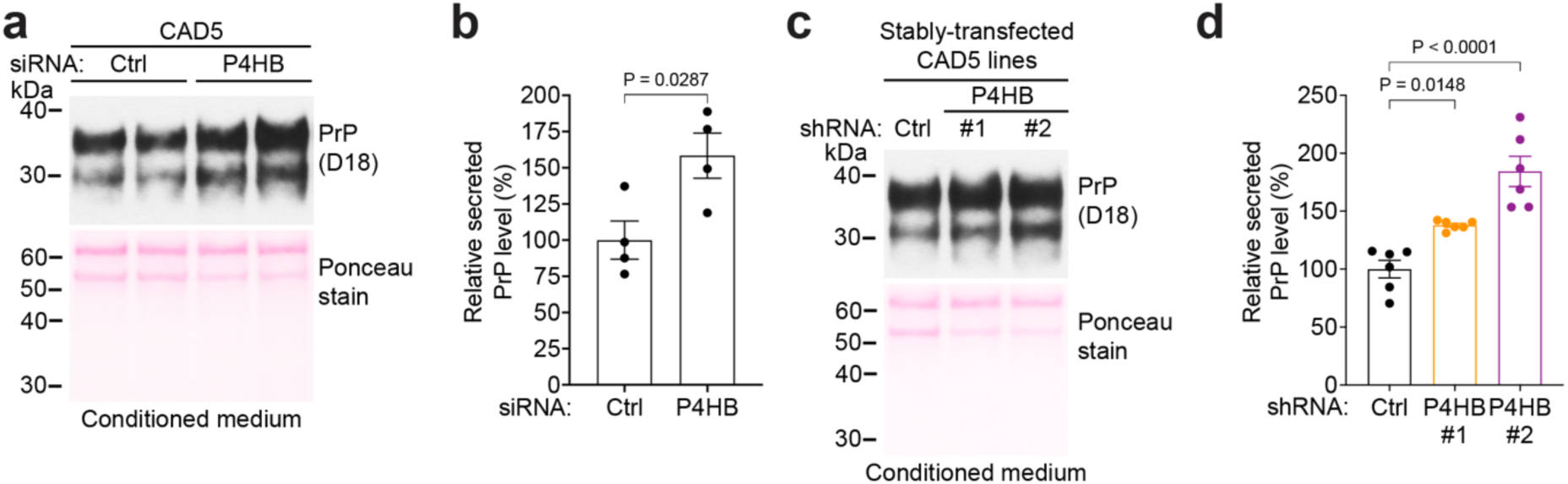
Knock-down of P4HB increases PrP secretion. (**a**) Immunoblot of secreted PrP levels in conditioned medium from CAD5 cells transfected with control or P4HB siRNA. The membrane was stained for total protein following transfer using Ponceau S. (**b**) Quantification of secreted PrP levels in CAD5 cells transfected with control or P4HB siRNA (n = 4 independent replicates). Data is mean ± SEM and statistical significance was assessed using an unpaired two-tailed t test. (**c**) Immunoblot of secreted PrP levels in conditioned medium from CAD5 cells stably transfected with control or one of two distinct P4HB shRNAs. The membrane was stained for total protein following transfer using Ponceau S. (**d**) Quantification of secreted PrP levels in CAD5 cells stably transfected with control or P4HB shRNA (n = 6 independent replicates). Data is mean ± SEM and statistical significance was assessed using one-way ANOVA followed by Dunnett’s multiple comparisons test.

**Figure S3.**
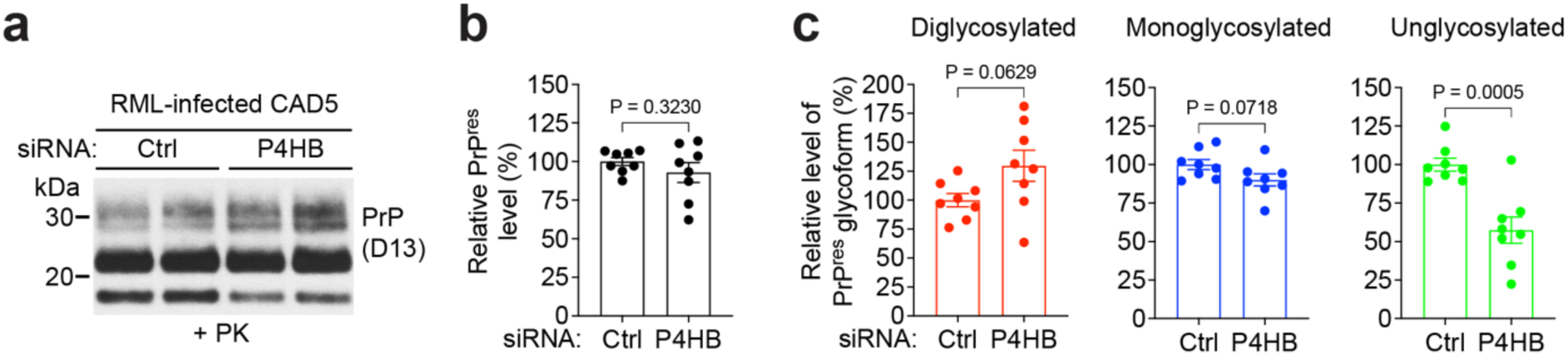
Knock-down of P4HB alters PrP^res^ glycosylation in RML prion-infected CAD5 cells. (**a**) Immunoblot of proteinase K (PK)-resistant PrP (PrP^res^) levels in RML prion-infected CAD5 cells transfected with control or P4HB siRNA for 72 h. (**b**) Quantification of PrP^res^ levels in RML prion-infected CAD5 cells transfected with control or P4HB siRNA (n = 8 independent replicates). Data are mean ± SEM and statistical significance was assessed using an unpaired two-tailed t test. (**c**) Quantification of PrP^res^ glycoforms in RML prion-infected CAD5 cells transfected with control or P4HB siRNA (n = 8 independent replicates). Data are mean ± SEM and statistical significance was assessed using an unpaired two-tailed t tests.

**Figure S4.**
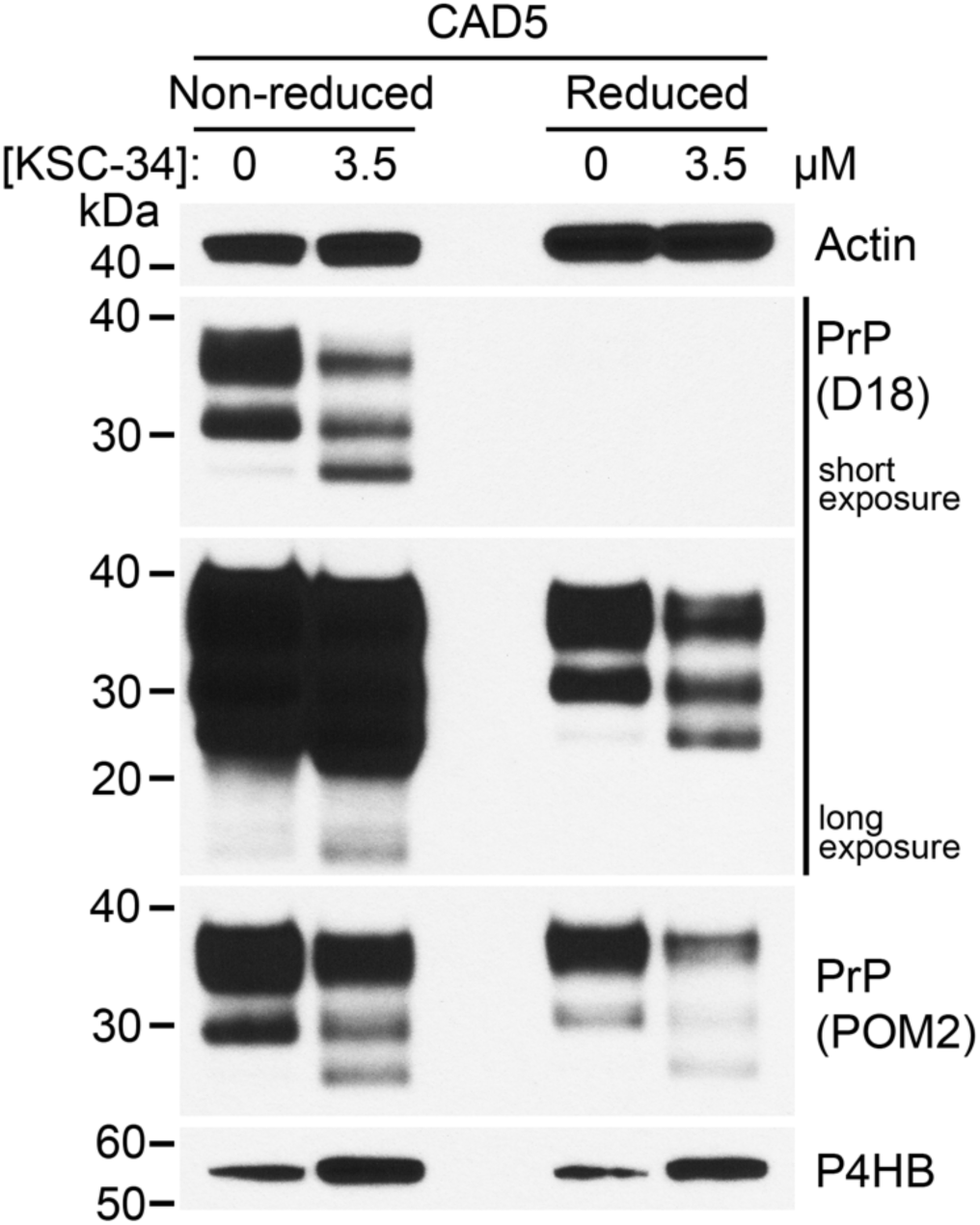
Analysis of PrP^C^ levels following P4HB inhibition using both non-reducing and reducing conditions as well as a second anti-PrP antibody. Representative immunoblots for PrP^C^ and P4HB in cell lysates from CAD5 cells treated with 0 or 3.5 μM KSC-34 for 72 h. In reduced conditions, lysates were treated with β-mercaptoethanol and boiled. PrP^C^ was detected using the antibodies D18 or POM2.

**Figure S5.**
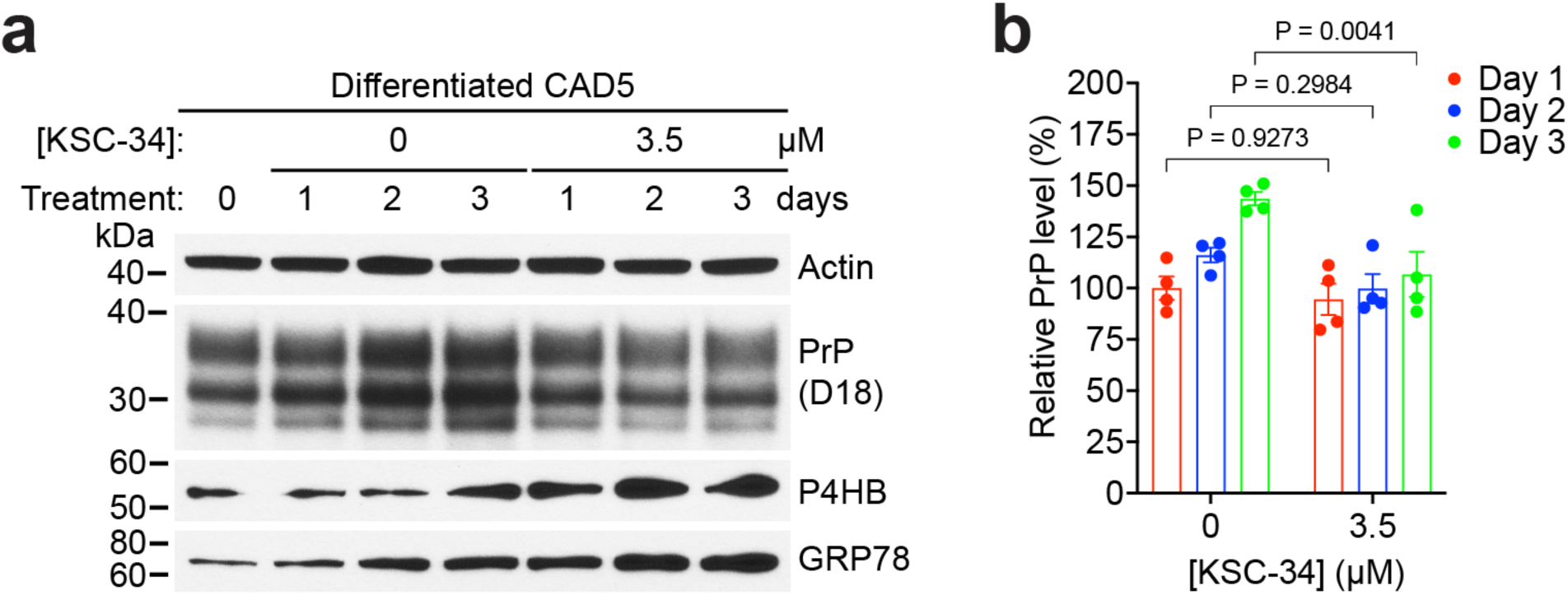
Inhibition of P4HB in differentiated (non-dividing) CAD5 cells decreases PrP^C^ levels. (**a**) Representative immunoblots of PrP^C^, P4HB, GRP78, and actin levels in differentiated CAD5 cells treated with 0 or 3.5 µM KSC-34 for 1, 2, or 3 days. (**b**) Quantification of PrP^C^ levels in differentiated CAD5 cells treated with 0 or 3.5 µM KSC-34 for 1, 2, or 3 days. The graph displays mean ± SEM and statistical significance was assessed using two-way ANOVA followed by Šídák’s multiple comparison test.

**Figure S6.**
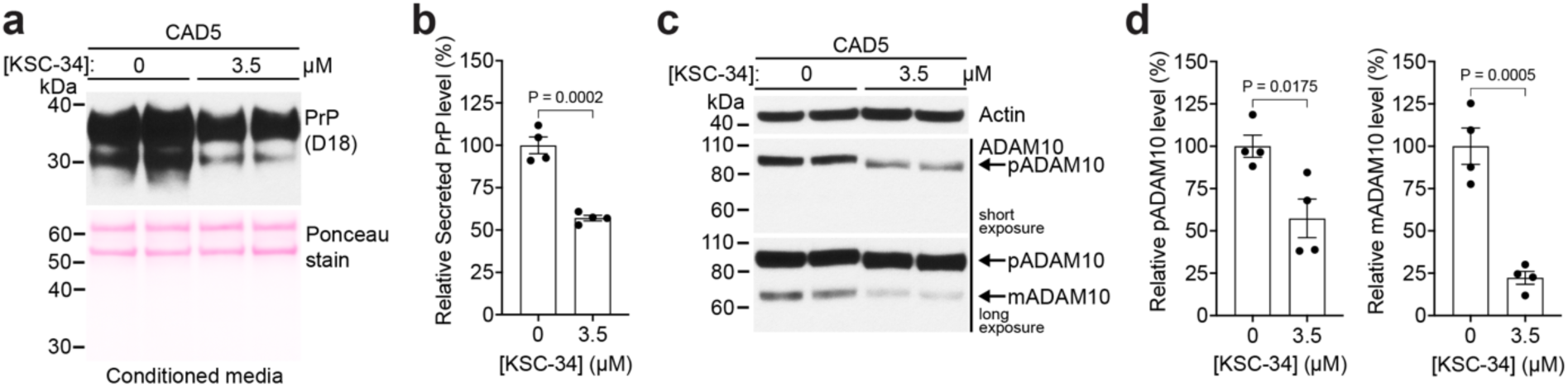
Inhibition of P4HB in CAD5 cells decreases levels of secreted PrP and ADAM10. (**a**) Immunoblot of secreted PrP levels in conditioned medium from CAD5 cells treated with 0 or 3.5 µM KSC-34 for 72 h. The membrane was stained for total protein following transfer using Ponceau S. (**b**) Quantification of secreted PrP levels in CAD5 cells treated with 0 or 3.5 µM KSC-34 (n = 4 independent replicates). Data is mean ± SEM and statistical significance was assessed using an unpaired two-tailed t test. (**c**) Immunoblots of the precursor (p) and mature (m) forms of ADAM10 in cell lysates from CAD5 cells treated with 0 or 3.5 µM KSC-34 for 72 h. Cell lysates were treated with β-mercaptoethanol and boiled prior to immunoblotting. (**d**) Quantification of pADAM10 and mADAM10 levels in lysates from CAD5 cells treated with 0 or 3.5 µM KSC-34 (n = 4 independent replicates). Data is mean ± SEM and statistical significance was assessed using unpaired two-tailed t tests.

**Figure S7.**
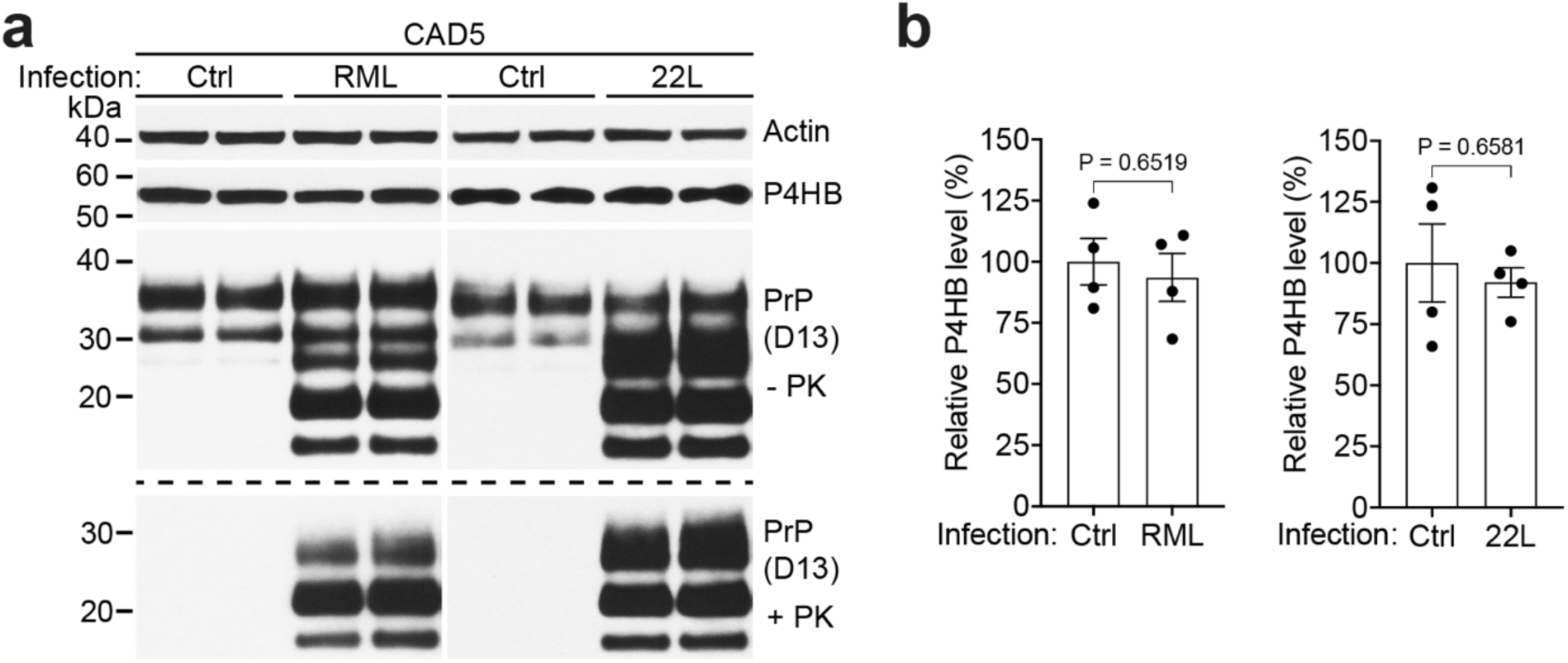
Prion infection does not modulate P4HB levels in CAD5 cells. (**a**) Representative immunoblots of total PrP (-PK), PrP^res^ (+PK), P4HB, and actin levels in cell lysates from mock-infected (Ctrl) CAD5 cells and CAD5 cells that have been stably infected with either RML or 22L prions. (**b**) Quantification of P4HB levels in cell lysates from Ctrl-infected CAD5 cells and CAD5 cells that have been stably infected with either RML or 22L prions. Data is mean ± SEM from 4 independent replicates. Statistical significance was assessed using unpaired two-tailed t tests.

**Figure S8.**
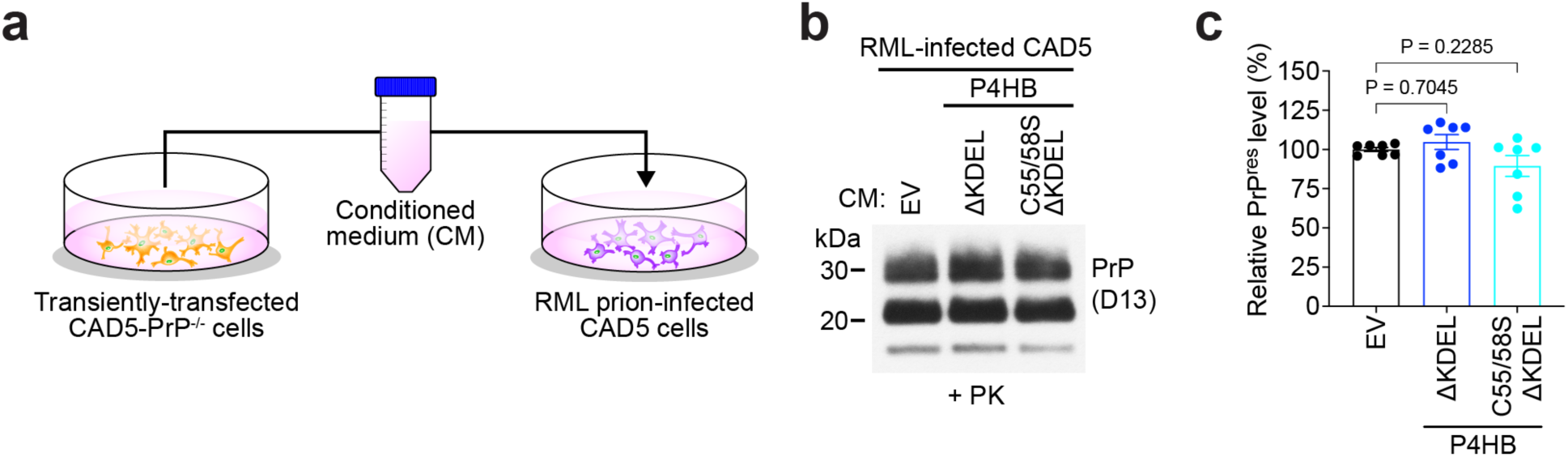
Conditioned medium containing P4HB does not alter PrP^res^ levels in RML prion-infected CAD5 cells. (**a**) Schematic of the experiment in which conditioned medium (CM) from transiently transfected CAD5-PrP^-/-^ cells is applied to RML prion-infected CAD5 cells for 72 h. (**b**) Representative immunoblot of proteinase K (PK)-resistant PrP^Sc^ (PrP^res^) levels in lysates from RML prion-infected CAD5 cells treated with CM from CAD5-PrP^-/-^ cells transiently transfected with empty vector (EV) or secreted P4HB variants. (**d**) Quantification of PrP^res^ levels in lysates from RML prion-infected CAD5 cells treated with CM from CAD5-PrP^-/-^ cells transiently transfected with EV or secreted P4HB variants (n = 7 independent replicates). The graph displays mean ± SEM and statistical significance was assessed using one-way ANOVA followed by Dunnett’s multiple comparisons test.

